# A SWI/SNF-specific Ig-like domain, SWIFT, is a transcription factor binding platform

**DOI:** 10.1101/2025.08.01.667725

**Authors:** Siddhant U. Jain, Kaylyn E. Williamson, Alexander W. Ying, Aasha M. Turner, Ruidong Jerry Jiang, Shaunak Raval, Kevin So, Maxwell J. Allison, Akshay Sankar, Daniel D. Sáme Guerra, Yutong Lin, Zhe Jiang, Nazar Mashtalir, Henry W. Rohrs, Cheryl F. Lichti, Tom W. Muir, Malvina Papanastasiou, Joao A. Paulo, Steven P. Gygi, Michael L. Gross, Cigall Kadoch

**Affiliations:** Department of Pediatric Oncology, Dana-Farber Cancer Institute and Harvard Medical School, Boston, MA 02215, USA; Broad Institute of MIT and Harvard, Cambridge, MA 02142, USA; Department of Chemistry, Washington University in St. Louis, St. Louis, MO 63105, USA; Department of Chemistry, Princeton University, Princeton, NJ 08540, USA; Departments of Pathology and Immunology, School of Medicine, Washington University in St. Louis, St. Louis, MO 63110, USA; Department of Cell Biology, Harvard Medical School, Boston, MA 02115, USA; Howard Hughes Medical Institute, Chevy Chase, MD 20815, USA

## Abstract

Mammalian SWI/SNF (BAF) chromatin remodeling complexes modulate DNA accessibility and gene expression, however, their genomic targeting mechanisms remain incompletely understood. Here, we identify SWIFT (SWI/SNF Ig-Fold for Transcription Factor Interactions), a conserved, broad transcription factor (TF) binding platform on the SMARCD subunits. SWIFT is necessary and sufficient for direct engagement with the transactivation domain of PU.1, a single mutation in which disrupts PU.1-mSWI/SNF binding, impairs complex targeting, and attenuates oncogenic transcription and proliferation in PU.1-dependent cancer cells. Dominant expression of the SWIFT domain in isolation sequesters TFs from mSWI/SNF and poisons TF-addicted cancer cells. Finally, TFs across diverse families interact with SMARCD paralog-specific SWIFT domains. These results define a major mechanism of cell type- and disease-specific mSWI/SNF chromatin targeting and inform approaches toward therapeutic modulation.

## Main Text

Timely and appropriate eukaryotic transcription requires highly coordinated, dynamic interactions among protein factors and DNA. Interplay between specialized proteins such as TFs, chromatin landscape features such as histone and DNA post-translational modifications, and regulatory machines such as chromatin remodeling complexes work in concert to control chromatin accessibility and binding of transcriptional apparatuses to target genes (*1*). Specifically, the mammalian <u>Swi</u>tch/<u>S</u>ucrose <u>N</u>on-<u>F</u>ermenting (mSWI/SNF) family of chromatin remodeling complexes encompasses a heterogenous collection of 11-15-subunit entities that use ATP hydrolysis to alter DNA-nucleosome contacts and hence remodel chromatin architecture (*2–7*). The mSWI/SNF complexes are found in three major configurations, termed canonical BAF (cBAF or BAF), polybromo-associated BAF (PBAF), and non-canonical BAF (ncBAF), each of which incorporate specific subunits and exhibit distinct localization proclivities and nucleosome interactions on chromatin (*8–16*). Within each complex, several subunit positions can be occupied by different subunits of a given paralog family (i.e. ARID1A/B, SMARCA4/2, SMARCD1/2/3, others) that are often co-expressed and assembled into complexes in a mutually exclusive manner. The resulting hundreds of specialized combinatorial mSWI/SNF assemblies play critical roles in development, cell type maintenance, and differentiation across diverse tissue types (*17–29*).

Notably, genes encoding mSWI/SNF complexes are mutated in over 20% of human cancers, making these remodelers the most frequently perturbed cellular entities second only to TP53 (*3*, *30*, *31*). Further, many cancers display vulnerabilities to mSWI/SNF gene disruption, including those with mSWI/SNF mutations which most often show enhanced sensitivity to paralog depletion (i.e. ARID1B in ARID1A-mut cancers, SMARCA2 in SMARCA4-mut cancers) or residual complex activities (i.e. ncBAF dependency in cBAF-perturbed cancers) (*8*, *10*, *15*, *32–37*) as well as transcriptionally addicted cancers lacking mSWI/SNF lesions (i.e. POU2F3-driven SCLC) (*38–42*). Genome-scale CRISPR-based fitness screens reveal that mSWI/SNF complex genes show strong co-dependencies with TFs, indicating their shared pro-oncogenic roles; conversely, cancer phenotypes can arise from cell lines bearing mSWI/SNF subunit mutations that stifle DNA accessibility generation over TF-controlled, lineage-specific enhancers that are required for proper differentiation (*38*, *43*, *44*). Taken together, these findings implicate mSWI/SNF complexes as critical determinants of TF-mediated gene expression.

Notably, mSWI/SNF subunits do not contain sequence-specific DNA-binding domains and contain only a limited set of histone reader domains, leaving open the question as to how mSWI/SNF complexes are differentially targeted to specialized sites genome-wide. While we determined histone landscape features that impact mSWI/SNF family complex binding and nucleosome remodeling activities (*16*), these findings are insufficient in isolation to explain the highly distinct mSWI/SNF complex targeting profiles observed across diverse human cell and tissue types. Further, several studies have identified indirect protein-level interactions between TFs and mSWI/SNF complexes, including those that are overexpressed or are part of fusion proteins in cancers as well as in models of normal cell development and differentiation (*21*, *22*, *45–57*). However, whether these interactions are direct and which subunits or region(s) within multicomponent mSWI/SNF complexes mediate potential TF interactions have not been identified to date. Defining the biochemical mechanisms by which TFs spanning different families and functional groups guide the targeting and activity of this major class of chromatin remodeling complexes is critical to understanding TF-addicted cancers, informing the determinants of mammalian developmental and differentiation processes, and to informing new strategies for therapeutic intervention.

## Results

### Transcription factor expression and cognate motif density directs tissue-specific genome-wide mSWI/SNF complex localization

Examination of mSWI/SNF complex occupancy across diverse cell types demonstrates enrichment of TSS-distal DNA motifs corresponding to one or more key lineage-specific transcription factors (TFs) (**fig. S1A-G**). Generally, mSWI/SNF genomic occupancy runs as a function of TF mRNA expression in a given tissue type and site-specific occupancy localized near TF target motifs (**fig. S1H-L**). There are over 30 different families of human transcription factors (TFs) as characterized by the structure of their DNA binding domains (*58*). To examine the impact of TF expression on mSWI/SNF complex occupancy and resulting accessibility, we individually expressed a diverse collection of HA-tagged TFs (n=14) spanning 10 TF families in an isogenic setting of human mesenchymal stem cells (MSCs), chosen for their plasticity(*59*) and profiled resulting TF and mSWI/SNF complex occupancy (CUT&RUN), chromatin accessibility (ATAC-seq) and transcriptional activity (RNA-seq) (**Fig. 1A and fig. S2A-B**).

**Figure 1.**
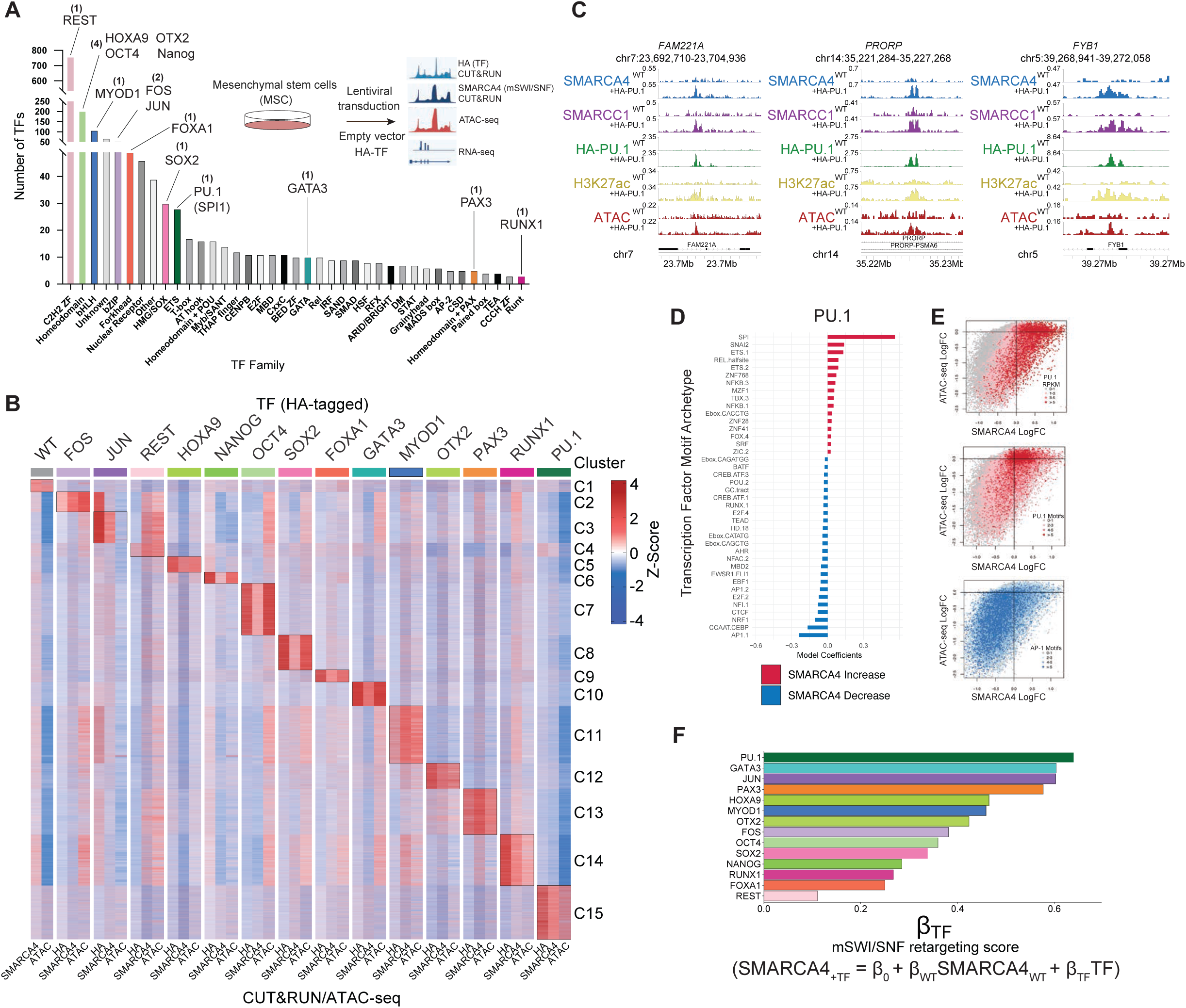
Transcription factors direct mSWI/SNF complex targeting and activity genome-wide. **(A)** Bar graph depicting human transcription factors grouped by family; those selected for HA-tagged expression and genomic studies in hMSCs are indicated. **(B)** Heatmap displaying the z-score normalized RPKM occupancies of HA-tagged TFs (as shown on top in colored boxes), SMARCA4, and DNA accessibility (ATAC-Seq) at all merged SMARCA4 and ATAC-Seq sites across all samples. Z-scores were calculated for each experiment individually prior to grouping of columns by cell lines for data visualization. Unguided hierarchical clustering was performed on using z-score normalized values for HA-TF CUT&RUN, which identified 18 distinct clusters as shown on the right. **(C)** Representative site showing TF-dependent mSWI/SNF localization and accessibility at the *FAM221A, PRORP,* and *FYB1* loci. RPKM-normalized enrichment of HA-tagged PU.1, SMARCA4, H3K27ac, DNA accessibility (ATAC-Seq) are shown. **(D)** GLMnet motif enrichment analysis was performed to identify top TF motifs underlying SMARCA4 peaks that display gain of SMARCA4 enrichment (in red) and loss of enrichment (in blue) upon PU.1 overexpression. **(E)** Scatterplots displaying the correlation between change in SMARCA4 occupancy (x-axis) and DNA accessibility measured by ATAC-Seq (Y-axis) in MSCs expressing PU.1 compared to empty vector. Color key indicates PU.1 RPKM enrichment (top panel), number of PU.1-motifs (middle panel) and number of AP.1 motifs (lower panel). **(F)** Bar chart displaying the standardized coefficient (βTF) for indicated HA-tagged TF from the linear regression model shown below.

Notably, overexpression of all TFs evaluated, including PU.1 (encoded by *SPI1* gene and henceforth, referred to as PU.1 for simplicity), GATA3, and others led to de novo gains in mSWI/SNF complex occupancy and DNA accessibility at highly TF-specific sites as measured by SMARCA4 CUT&RUN and ATAC-seq peak signals, respectively, at previously mSWI/SNF-deficient and inaccessible genomic loci (**Fig. 1B-C**, and fig. **S2C**). Strikingly, TF overexpression in hMSCs resulted in TF-specific sets of sites of mSWI/SNF targeting genome-wide at which accessibility was generated, and nearby gene expression was increased (**Fig. 1B, and fig. S2D).** Expression of REST was an exception, in line with its role in transcriptional repression (*60–62*). These results are exemplified at selected target sites shown for PU.1, GATA3, and PAX3 factors (**Fig. 1C, and fig. S2D-F**).

Notably, TF expression resulted in skewing of mSWI/SNF occupancy to sites enriched for the archetypal motif of the primary overexpressed TF, coupled with concomitant depletion of occupancy over AP-1 family motifs, as visualized by an unbiased linear regression model (GLMnet) of the fold change TF motif archetype enrichment under SMARCA4 target sites (**Fig. 1D, and fig. S2G-H**). This is exemplified in the setting of PU.1 and GATA3 expression which led to SMARCA4-marked mSWI/SNF occupancy showing top enrichment for the corresponding SPI and GATA motifs, respectively, with reductions in occupancy over AP-1 and CEBP motif-containing sites (**Fig. 1D, and fig. S2H**). This supports a potential model in which tissue-specific TFs dominate, either by expression (abundance), affinity, or a combination thereof, over general mSWI/SNF-interacting TFs such as AP.1 in directing mSWI/SNF complex localization genome-wide. Notably, sites of greatest increases in mSWI/SNF complex occupancy and accessibility were those enriched with the greatest number of motifs corresponding to the overexpressed TF (**Fig. 1E, and fig. S2I-J**). These sites are also those most highly bound by the expressed TF itself, suggesting motif-based concentration dependent recruitment of the TF and mSWI/SNF complexes to de novo sites (**Fig. 1E**, **and fig. S2I**). This is demonstrated at the single locus level at distal enhancer sites with 5, 3, and 1 PU.1 (PU.1) archetypal motifs nearest the *A2M, HGD,* and *CD48* loci, respectively, to which PU.1 itself, as well as mSWI/SNF complexes are recruited in a motif-dependent manner resulting in a gradient of changes in accessibility and nearest gene expression (**fig. S2K**). Sites with AP-1 family TF consensus motifs were depleted for mSWI/SNF complex occupancy and chromatin accessibility upon expression of lineage-specific TFs, further corroborating the depletion from these sites observed in the unbiased linear regression model (**Fig. 1E, and fig. S2H**). Taken together, these results indicate a TF-specific, TF motif concentration-dependent impact on the genome-wide occupancy of mSWI/SNF complexes and their activities.

### Mass-spectrometry-based footprinting resolves a PU.1-mSWI/SNF interaction region in the structural core module

Based on these findings, we next sought to determine whether TFs and mSWI/SNF complexes directly interact. We purified human PU.1, a TF that showed the highest correlation with de novo SMARCA4 targeting in cells, as well as fully assembled cBAF complexes from mammalian cells (HEK-293T) using overexpression and subsequent density gradient sedimentation (**Fig. 2A, and fig. S3A-B**)(*9*, *11*). Indeed, we incubated resultant proteins in vitro and found that PU.1 interacts directly with cBAF complexes in the absence of any additional factors (**Fig. 2B**). PU.1 contains an N-terminal disordered region containing an acidic and glutamine-rich domain (A/Q-rich, aa 10-100) and a PEST (proline/glutamate/serine/threonine-rich) domain. Based on the sequence compositional similarity with transactivation domains (tAD) of TFs (*63–67*), we refer to this disordered region (aa 10-160) as the tAD henceforth. Notably, deletion of the PU.1 disordered tAD (residues 10-160) nearly completely abrogated interaction with purified cBAF complexes (**Fig. 2B**). In line with these in vitro results, overexpression of a PU.1 ΔtAD deletion mutant in hMSCs failed to recruit mSWI/SNF complexes, marked by SMARCA4, to PU.1 target sites, relative to full-length PU.1, suggesting that the tAD region is necessary both for direct biochemical interaction and PU.1-mediated mSWI/SNF targeting (**Fig. 2C)**.

**Figure 2.**
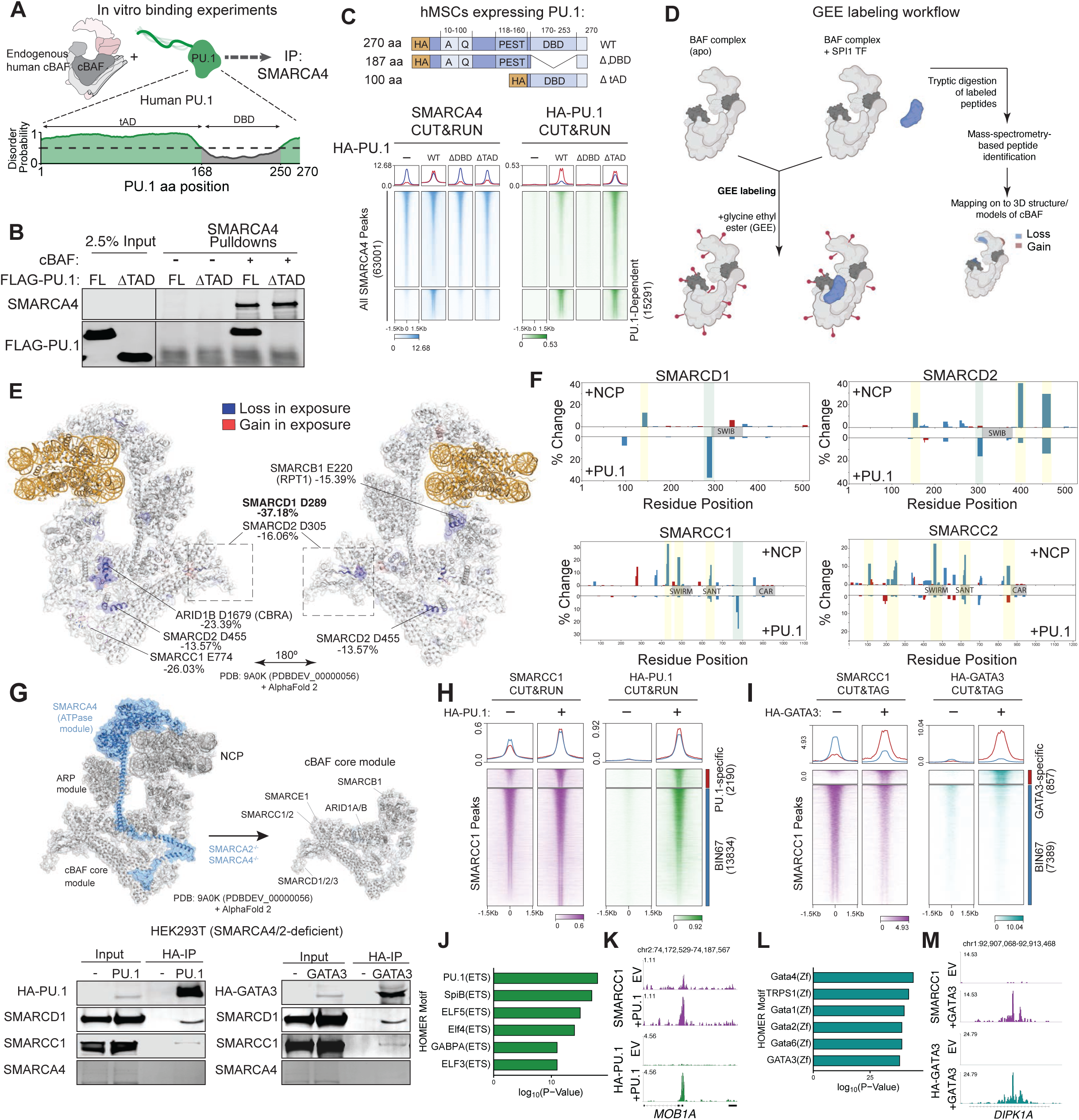
PU.1 interacts with the mSWI/SNF core structural module. **(A)** Schematic for in vitro incubation experiments using endogenous, fully assembled human cBAF complexes and full-length human PU.1. Prediction of disordered domains within PU.1 within tAD and DBD are indicated below. **(B)** SMARCA4 pulldown experiments performed with 2.5 µg of purified cBAF incubated with 10-fold molar excess of PU.1 wild-type or PU.1 ΔtAD (Δ1-160). **(C)** (Top) schematic for HA-tagged PU.1 variants introduced into human MSCs for PU.1 and SMARCA4 ChIP-seq experiments; (Bottom) heatmaps depicting PU.1 and SMARCA4 occupancy across all merged SMARCA4 sites in empty vector, and PU.1 WT, delDBD, and delTAD variant conditions. PU.1-specific target sites are indicated. **(D)** Schematic for mass-spec-based GEE protein footprinting experiments performed with endogenous human cBAF complexes and full-length PU.1. **(E)** cBAF peptides displaying GEE-labeling changes were mapped onto the 3D structure of NCP-bound cBAF, with SMARCD2 modeled with AlphaFold2 and superimposed on PDB:9A0K (PDBDEV_00000056). **(F)** Bar charts across mSWI/SNF SMARCD and SMARCC subunits depicting %GEE labeling change upon incubation with NCP (top) or PU.1 (bottom). **(G)** (Top), schematic depicting mSWI/SNF complexes with a stable and independently assembled core upon removal of the ATPase subunits; (bottom) Expression of HA-tagged PU.1 or GATA3 TFs in SMARCA4/SMARCA2 dual-deficient HEK-293T cells results in mSWI/SNF complex interactions. **(H-I)** Heatmaps depicting occupancy of mSWI/SNF core (SMARCC1) with and without expression of HA-tagged PU.1 (**H**) or HA-tagged GATA3 (**I**). **(J-K)** HOMER motif analyses performed on SMARCC1 sites gained in the cells overexpressing PU.1 (**J**) or GATA3 (**K**). **(L-M)** Same as J and K but for a GATA3-dependent sites.

Next, we developed and implemented a biochemical approach to define the cBAF complex subunit or interface with which the PU.1 TF interacts. We used a chemical labeling method using water-soluble carbodiimde in combination with glycine ethyl ester (GEE) to specifically modify solvent-accessible carboxyl groups on glutamate (E) and aspartate (D) residues (*68–70*), thus reporting on changes in surface exposure upon PU.1-cBAF interaction in vitro (**Fig. 2D**). In parallel, we performed these studies with cBAF incubated with recombinant nucleosomes, a well-documented and structurally resolved interaction, as a control using the same conditions. Notably, we identified 75 unique changes in GEE labeling upon PU.1 binding, including 57 sites with decreased and 18 sites with increased surface exposure (**Fig. S3C-D**). GEE labeling changes detected on cBAF complexes upon NCP binding mapped largely to the helicase domain of the SMARCA2/SMARCA4 ATPase subunits and other structural core subunits that have been shown to undergo configurational rearrangement upon NCP engagement (**Fig. S3C-E)** (*9*, *71–73*). The top 7 sites with the most substantial reductions in surface exposure upon PU.1-cBAF incubation all mapped to subunits of the structural core module of mSWI/SNF, including SMARCD and SMARCC subunits (**Fig. 2E-F, and fig. S3D**). Residue D289 of the SMARCD subunits (shared between paralogs) exhibited the largest reduction in surface accessibility upon PU.1 incubation (-37.18% decrease relative to unbound). These data inform potential sites of direct interaction of the PU.1 TF with the cBAF complex core module and nominate the SMARCD1/2/3 core module subunits as those containing regions with putative direct interfaces.

We have previously demonstrated that a structural core module of cBAF complexes comprising of SMARCD1/2/3, SMARCC1/2, SMARCB1, SMARCE1 and ARID1/ARID2 subunits is formed in cells independent of the presence of the catalytic ATPase module containing SMARCA2/4 (**Fig. 2G**) subunits (*9*, *74*). Consistently, PU.1 and another lineage-specific TF, GATA3, both co-immunoprecipitated SMARCC1 and SMARCD1 subunits in HEK-293T cells genetically deficient in SMARCA2/4 subunits (**Fig. 2G, and fig. S4A**), indicating that the cBAF core is sufficient to interact with PU.1 and GATA3 TFs in cells. Further, in a cancer cell line deficient in SMARCA2/4 and hence lacking the entire ATPase module (BIN-67 SCCOHT cells)(*74–77*), we found that expression of PU.1 or GATA3 maintained ability to redirect core module occupancy on chromatin (**Fig. 2H-I, and fig. S4B-E**), with TF-specific de novo mSWI/SNF sites showing corresponding motif enrichment (**Fig. 2J-M**). As expected, reintroduction of WT but not ATPase-dead (K785R mutant) SMARCA4 into BIN-67 ATPase-deficient cells resulted in substantial increases in DNA accessibility (*74*) and additional mSWI/SNF target sites, including those targeted by PU.1 and GATA3 TFs (**fig. S4F-I**).

### An Ig-like fold, SWIFT, in the mSWI/SNF structural core interacts with PU.1 in vitro

The region on cBAF complexes that showed highest reduction in GEE labeling upon PU.1 incubation mapped to a functionally uncharacterized domain in the SMARCD subunits (**Fig. 2**). Encompassing the residues that displayed the greatest changes in surface exposure is a ∼26 KDa domain that contains a sandwich arrangement of 8 anti-parallel β-strands, characteristic of the immunoglobulin folds (Ig-fold) (**Fig. 3A, and fig. S5A-B**)(*78*, *79*). However, the SMARCD1/2 β-sandwich deviates from conventional Ig-folds by incorporating a SWIB domain within loop 7, which we and others have demonstrated interacts with the ARID1A/1B or ARID2 subunits and is necessary for complex assembly (**Fig. 3A, and fig. S5A**)(*9*, *72*, *80*). Sequence alignment of the SMARCD1/2 Ig-fold suggested that it is distantly related to the YEATS domains found in chromatin modifiers (AF9, ENL etc.) (**Fig. S5C**) (*81–84*). However, aromatic residues within loop 6 of the YEATS domain, which constitute an aromatic tunnel and are essential for binding histone acylation through ∏-∏-∏ stacking, are notably absent from the SMARCD1/2/3 Ig-fold, suggesting a divergent function (**Fig. 3A, and fig. S5C-D**). Indeed, the SMARCD2 Ig-fold domain failed to interact with H3K9crotonylated peptides in vitro, compared to AF9 which, as expected, displayed strong binding (**Fig. S5E-F)**. Due to its close proximity with nucleosomal DNA in the context of nucleosome-bound mSWI/SNF complexes, previous studies have hypothesized that the SMARCD YEATS-like domain may interact with TFs, however, its biochemical function has not been evaluated (*73*, *85*). Based on our characterization of this SMARCD Ig-fold, we will henceforth refer to it as **SW**I/SNF **<u>I</u>**<u>g</u>-**<u>F</u>**old for **<u>T</u>**F-interactions (SWIFT).

**Figure 3.**
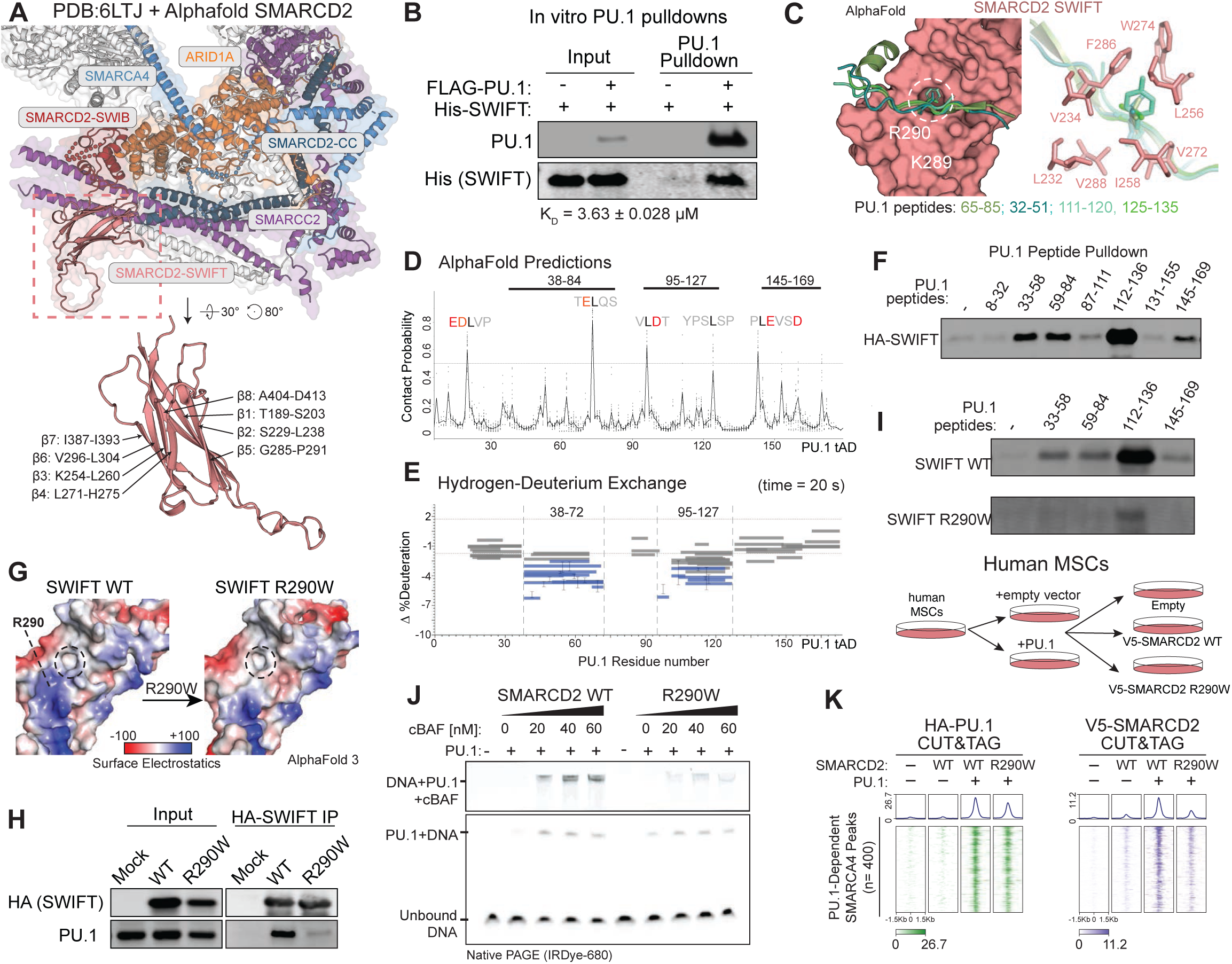
The SMARCD2 SWIFT domain is necessary and sufficient to for interaction with PU.1 in vitro and in cells. **(A**) (Top) mSWI/SNF core subunits showing interactions between SMARCD2 (salmon) coiled coil domain with SMARCC2 (blue), and between SWIB domain with ARID1A (orange); (Bottom) Structure of the SMARCD2 Ig-like fold, SWIFT; **PDB: 6LTJ**+ Alphafold SMARCD2. **(B)** Immunoblots of FLAG-PU.1 pulldown experiments performed by incubation of 0.5 µM FLAG-tagged PU.1 was incubated with 2 µM SMARCD2 SWIFT domain. **(C)** AlphaFold3 multimer modeling displaying the putative interaction between PU.1 20-mer peptides and a hydrophobic pocket on the surface of SMARCD2 SWIFT domain. **(D)** Contact probability scores of interactions between each residue on the transactivation domain of PU.1 with the residues within the hydrophobic pocket on SWIFT domain determined by Alphafold modeling of all consecutive PU.1 10-mer peptides. Line shows the mean of all outcomes for a given residue. Solid line on top marks the regions that showed binding to the SWIFT domain in vitro in **E** and **F**. **(E)** Change in % Deuterium uptake by each PU.1 peptide upon addition of SMARCD2 SWIFT domain. Dashed line represent the threshold of statistically significant changes. Error bars represent standard deviation. **(F)** Immunoblots of eluates from peptide pulldown of 7 PU.1 peptides (25-mer) spanning full transactivation (tAD) domain. **(G)** 3D-surface rendering of surface electrostatic potential of wildtype SMARCD2 SWIFT and R290W point mutation near the hydrophobic pocket (shown in black dashed circle). **(H)** Immunoblots displaying the eluates from Immunoprecipitation of HA-tagged SWIFT domain from HEK-293T cells expressing HA-tagged SWIFT wildtype or R290W mutant and FLAG-tagged PU.1. **(I)** Immunoblots of eluates from peptide pulldowns of PU.1 peptides with HA-tagged SMARCD2 SWIFT wildtype (top) or R290W mutant (bottom). **(J)** Electromobility shift assays (EMSA) performed on complex formed by dsDNA containing a PU.1 binding motif and PU.1 incubated with cBAF complexes containing wildtype SMARCD2 or R290W mutation. Upper panel shows the supershift caused by addition of cBAF complexes. **(K)** Heatmaps depicting PU.1 and V5-tagged SMARCD2 occupancy (CUT&RUN) in hMSCs expressing empty vector, V5-tagged WT SMARCD2 or SMARCD2 R290W mutant in the presence or absence of PU.1.

We incubated purified SMARCD2 SWIFT domain and full-length PU.1 in vitro (**Fig. S5F**). Remarkably, SMARCD2 SWIFT displayed direct interaction with purified PU.1 outside of the context of fully assembled mSWI/SNF complexes in in vitro pulldown experiments (**Fig. 3B**). Using microscale thermophoresis (MST), we determined the dissociation constant for the interaction between the SMARCD2 SWIFT domain and PU.1 as 3.63 µM (**Fig. S6A**), which is comparable to those measured for TF-P300 interactions and other protein-protein interactions (*65*, *84*, *86*, *87*). These data establish that the SMARCD2 SWIFT domain is sufficient for interaction with PU.1 in vitro.

Next, we modeled the SMARCD2 SWIFT and PU.1 interaction using Alphafold, which determined interaction between PU.1 residue L73 within its tAD domain with a hydrophobic pocket formed between the two sandwiched β-sheets of the SWIFT domain (F286, W274, V288, V272) with an aggregate contact probability of 81% and lowest predicted alignment error of 4.6 (**fig. S6B**). The adjacent residue PU.1 F71 interacted with SWIFT R233 through cation-pi interactions. Moreover, PU.1 aa 80-95 formed a transient ɑ-helix with SWIFT V257 and L271 in which PU.1 L81 and L84 engaged in hydrophobic interactions. However, mutations in these residues on PU.1 – F71A and L73A, as well as L84P to prohibit formation of the transient ɑ-helix – failed to impair PU.1-mediated genomic recruitment of mSWI/SNF in hMSCs (**fig. S6C-D**). These results suggest that such a singular monovalent interaction is insufficient to explain PU.1-mSWI/SNF interactions in cells.

TF tADs are not conserved evolutionarily or between families; however, these intrinsically disordered regions contain Short Linear Motifs (SLiMs) that are collectively enriched in hydrophobic and acidic residues, which are altogether necessary for transactivation (*63*, *65–67*, *88*). These features led us to hypothesize that SLiMs in the PU.1 tAD make allovalent interactions with the hydrophobic pocket on the SWIFT domain (*89–91*). In silico interaction prediction between the SWIFT domain and all consecutive 10-mer peptides in the PU.1 tAD identified that multiple putative PU.1 residues engage with the same hydrophobic pocket on the SWIFT domain (**Fig. 3C-D**). Due to the limitations of AlphaFold multimer models to predict protein-protein interactions, specially involving intrinsically disordered regions such as tADs, we sought to experimentally identify the SWIFT domain binding sites on PU.1 tAD using hydrogen-deuterium exchange (HDX) experiments by incubating the PU.1 tAD (aa 1-165) with an excess of SMARCD2 SWIFT. We identified two distinct regions of the PU.1 tAD – residues 38-72 and 95-127 – that showed equivalent binding with the SWIFT domain (**Fig. 3E** and **fig. S6E**), which may engage in either independent and equal binding events or represent a primary binding event, followed by a secondary avid interaction. To delineate, we synthesized seven 25-aa long peptides spanning the entire PU.1 tAD domain and tested their ability to independently bind the SWIFT domain (**Fig. 3F**). We found that 4 of the 7 peptides bound SMARCD2 SWIFT domain with varying levels, regions of which corroborated HDX findings. Together, these data suggest that the PU.1 tAD contains multiple sites able to engage with the SWIFT domain in an allovalent manner (**Fig. 3C-F**).

By in silico structural modeling, we identified acidic (Asp/Glu) or electronegative residues adjacent to nearly all tAD residues that interact with the SWIFT the hydrophobic pocket. Intriguingly, the surface adjacent to the SWIFT hydrophobic pocket contains several solvent-exposed positively charged residues, in particular, R233, K289 and R290 residues (**Fig. 3D, and fig. S6F**). We thus hypothesized that the PU.1 tAD makes allovalent interactions with SWIFT for which aliphatic and aromatic residues on the tAD engage with the hydrophobic pocket and that the adjacent amino acids make electrostatic or hydrogen bonds with the basic region. Consistent with this hypothesis, mutation of basic residues, including R290A, R290E and a cancer-associated missense mutation, R290W (COSMIC dataset), markedly reduced PU.1 binding to SMARCD2 SWIFT in cells (**Fig. 3G-H** and **fig. S6F-H**). Double- and triple-mutant combinations led to a graded reduction in PU.1 binding, indicating that each residue individually contributes to binding affinity. Notably, all PU.1 peptides lost interaction with the R290W mutant SWIFT domain in vitro (**fig. S6F-G**). The loss of interactions of all PU.1 peptides by a single point mutation near the hydrophobic cage suggest that SLiMs on PU.1 tAD engage in allovalent interactions with the SWIFT domain, distinct from a multivalent model of interaction (**Fig. 3I** and **fig. S6I**).

Finally, to characterize the PU.1-mSWI/SNF interaction through the SMARCD2 SWIFT, we purified fully assembled mSWI/SNF complexes containing either WT or R290W mutant variant SMARCD2 (**Fig. S6C**). As expected from the solvent-exposed R290 side chain within the SWIFT domain structure, R290 mutation did not affect mSWI/SNF complex integrity or remodeling activities (**fig. S7A-E**). However, consistent with impaired interaction with PU.1, mSWI/SNF complexes containing SMARCD2 R290W failed to supershift PU.1-DNA complexes in vitro (**Fig. 3J** and **fig. S7F-H**). Consistent with our *in vitro* results, expression of PU.1 in hMSCs resulted in recruitment of V5-tagged wild-type SMARCD2 to PU.1 target sites but markedly impaired recruitment of SMARCD2 R290W mutant-containing complexes (**fig. S7I**). Taken together, our results demonstrate that the SWIFT Ig-like domain is necessary and sufficient for the mSWI/SNF interaction with PU.1 in vitro and in cells.

### A single-residue mutation in SWIFT domain blocks PU.1-mediated mSWI/SNF targeting and impairs oncogenic gene expression and proliferation in transcription factor-driven AML cells

We next sought to determine whether alteration of the SMARCD2 SWIFT-PU.1 interaction is necessary to support the chromatin occupancy, gene regulatory and proliferative signatures of human cancers dependent on the PU.1 TF as a strategy to evaluate whether TF interactions with this domain may be required for TF-‘addicted’ cancers. To this end, PU.1 is central in orchestrating lineage commitment in the myeloid and macrophage lineage and is a top-ranked vulnerability in AML cell lines where it functions to maintain myeloid progenitor expression programs and self-renewal (*28*, *40*, *92–97*) (**fig. S8A**). Integrating results from genome-scale CRISPR-Cas9-based fitness screens performed across >1000 cancer cell lines, we find that AML cell lines exhibit among the highest co-dependency scores on both PU.1 and SMARCD2, suggesting the possibility that the interaction between PU.1 and SMARCD2 is necessary for oncogenic transcriptional functions in these cells (**Fig. 4A, and fig. S7A**). To assess this, we suppressed endogenous expression of SMARCD2 in MOLM13 cells (using shRNA) with simultaneous rescue of either WT SMARCD2 or SWIFT domain binding-deficient R290W mutant SMARCD2 using lentiviral overexpression (**Fig. 4B**). Consistent with our in vitro biochemical results and in-cell results, complexes containing the SMARCD2 R290W nearly completely ablated their interaction with endogenous PU.1 in AML cells, while maintaining interactions with other mSWI/SNF subunits (**Fig. 4C, and fig. S8B**).

**Figure 4.**
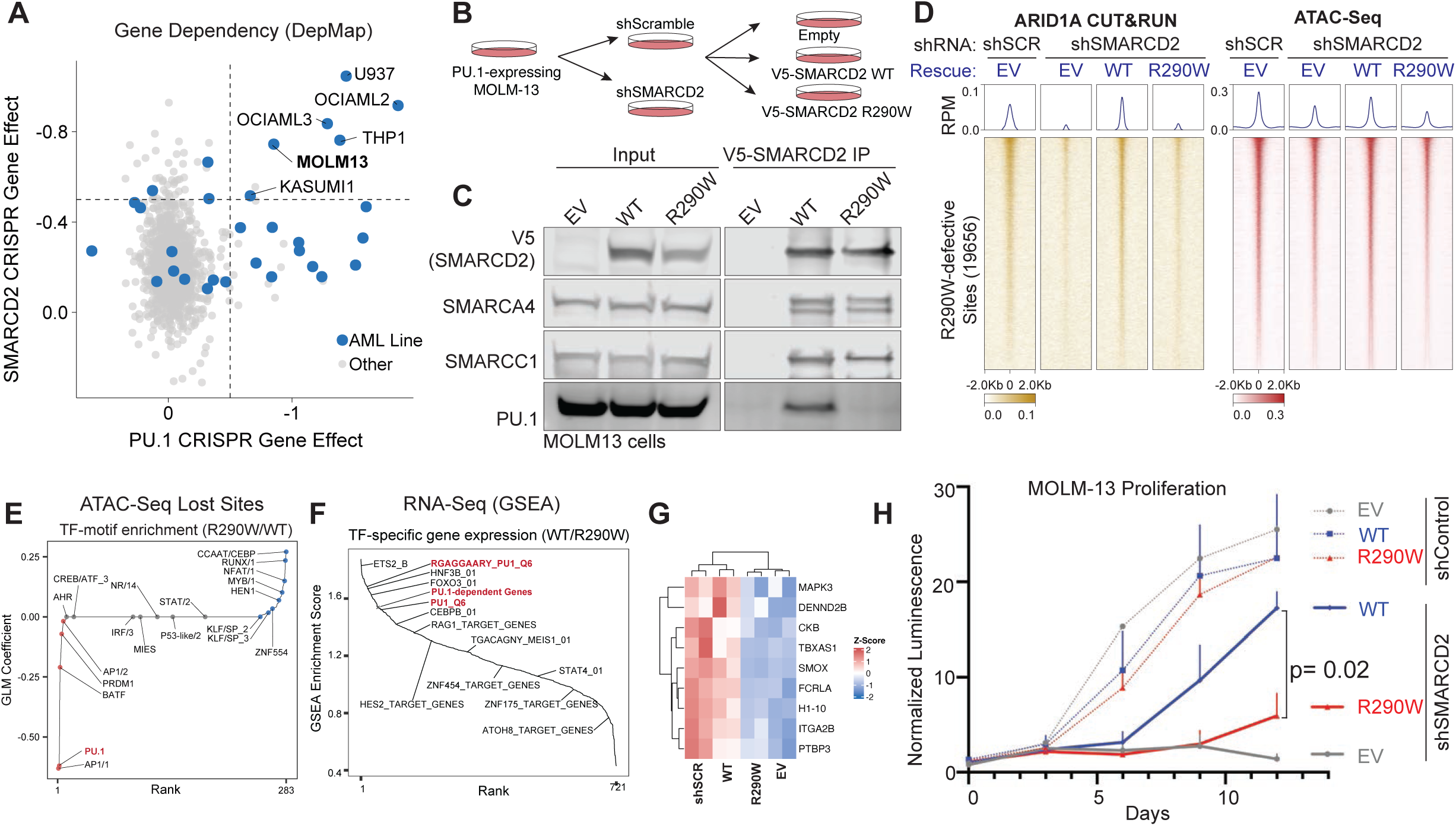
The SMARCD2 SWIFT domain is necessary to sustain PU.1-dependent transcription and cancer cell proliferation. (**A)** Scatterplot displaying CRISPR gene effect scores (dependency) for SMARCD2 (Y-axis) and PU.1 (X-axis) across n=900 cancer cell lines. Acute myeloid leukemia (AML) cell lines are indicated in blue. **(B)** Schematic displaying the experimental strategy to identify the function of PU.1-SWIFT interaction interface in MOLM-13 AML cells. Endogenous SMARCD2 is suppressed using shRNA followed by exogenous expression of shRNA-resistant SMARCD2 WT or R290W mutant variants. **(C)** Immunoblots performed on inputs and V5 immunoprecipitations of nuclear extracts isolated from MOLM-13 cells containing either WT or R290W mutant SMARCD2. (**D)** Heatmaps depicting ARID1A chromatin occupancy (CUT&RUN) and DNA accessibility (ATAC-seq) at cBAF-occupied sites in MOLM-13 cells rescued with empty vector (EV), WT SMARCD2, or R290W mutant following shRNA-mediated endogenous SMARCD2 suppression. (**E)** Motif enrichment analysis at genomic sites with reduced chromatin accessibility in cells rescued with the SMARCD2 R290W mutant. (**F)** Gene set enrichment analysis of genes downregulated in cells rescued with SMARCD2 R290W mutant compared to SMARCD2 WT. PU.1 target gene sets are highlighted in red. (**G)** Z-score normalized RNA-Seq expression of PU.1-target genes in MOLM-13 AML cells in control (shScramble) or shSMARCD2 rescued with either EV, WT SMARCD2 or R290W SMARCD2(*137*). (**H)** Cell proliferation (normalized luminescence) of MOLM13 cells with SMARCD2 knockdown (or a scrambled control) rescued with shRNA-resistant SMARCD2 WT or R290W transgenes. Proliferation was monitored for 12 days following infection and selection (data points represent 2 independent biologic replicates, each with 2 technical replicates).

We next aimed to define the impact of the SMARCD2 SWIFT domain mutation on the targeting and accessibility generation of mSWI/SNF complexes in human AML cells using CUT&RUN and ATAC-seq, respectively. As expected, loss of SMARCD2 resulted in decreased mSWI/SNF chromatin occupancy genome-wide, owing to its critical functions in initial mSWI/SNF core assembly (**fig. S8C-D**). Targeting of cBAF-specific subunit, ARID1A, was affected to a larger extent compared to pan-mSWI/SNF subunit, SMARCA4, consistent with previous studies demonstrating that SMARCD2 preferentially assembles into cBAF complexes, while ncBAF complexes exclusively incorporate the SMARCD1 paralog (**fig. S8E-J**). Importantly, compared to rescue expression of SMARCD2 WT, R290W mutant SMARCD2 failed to restore cBAF occupancy (ARID1A) globally at ∼80% sites, with the PU.1 motif among the top significantly enriched motifs underlying these peaks. Assessment of mSWI/SNF activity using ATAC-Seq identified a collection of 19,656 sites that required WT SMARCD2 for mSWI/SNF targeting in MOLM13 cells (**Fig. 4D, and fig. S8E-I**). Over these sites, targeting of mSWI/SNF complexes marked by both ARID1A and SMARCA4, as well as accessibility was markedly decreased (**Fig. 4D, and fig. S8F-G**). Genomic sites recovered by SMARCD2 R290W mutant likely represent SWIFT-independent mechanisms of mSWI/SNF recruitment, including through interactions with histone modifications or IDR-IDR interactions with other cofactors (*16*, *98*). Distance-to-TSS analyses revealed these sites largely over distal enhancers (**Fig S8K**); motif analysis performed on these sites identified PU.1 motif as the top motif affected (**Fig. 4E**). PU.1 occupancy was also reduced over these sites in the SMARCD2 KD and SMARCD2 R290W conditions, likely owing to reduced accessibility and open motif availability for binding (**Fig. S8L**).

Finally, we examined the impact of the R290W mutant on MOLM-13 AML gene expression using RNA-seq. Notably, GSEA analyses performed in SMARCD2 WT and R290W mutant rescue conditions revealed PU.1 target genes among top-scoring gene sets impacted by the single residue mutation (**Fig. 4F-G, and fig. S8M**). Key target genes important in the maintenance of AML leukemogenesis were rescued in expression following SMARCD2 knockdown only by WT SMARCD2 and not R290W mutant SMARCD2 (**Fig. S8I**). Finally, as MOLM-13 cells are dependent on PU.1 for proliferative maintenance (**Fig. 4A, and fig. S8A**), we sought to determine the impact of SMARCD2 loss and the ability of either WT SMARCD2 or mutant R290W SMARCD2 to rescue cancer cell line proliferation in culture. Strikingly, knockdown of SMARCD2 attenuated proliferation of MOLM-13 cells; rescue of this proliferative defect was achieved only with SMARCD2 WT and not with the R290W mutant (**Fig. 4H**). Together, these data substantiate our biochemical results by demonstrating that the SMARCD2 SWIFT domain is necessary for recruitment of mSWI/SNF by PU.1 in cells and the subsequent activity at PU.1 target genes necessary to uphold cancer cell proliferation.

### SWIFT domain is a mSWI/SNF-specific evolutionarily conserved TF binding platform

Our experiments thus far revealed that the SWIFT domain is necessary and sufficient for interaction with the PU.1 transcription factor in vitro and in cells. We next expressed PU.1 in hMSCs along with the SMARCD2 SWIFT domain (SMARCD2 aa185-415;Δ312-382) in isolation which is sufficient for interaction with PU.1 but cannot assemble into mSWI/SNF complexes due to the absence of its SWIB (aa 312-382) and coiled coil domains necessary for binding core subunits (*9*, *71–73*, *80*). Notably, while ectopic expression of PU.1 led to substantial de novo mSWI/SNF targeting, expression of PU.1 with concomitant expression of the SMARCD2 SWIFT markedly reduced complex occupancy at PU.1 target sites (**Fig. 5A, and fig. S9A-B**). Together, these data indicate that expression of the PU.1-interacting SWIFT domain in isolation blocks recruitment of endogenous mSWI/SNF at PU.1 target sites by dominantly interacting with, or “sequestering”, PU.1.

**Figure 5.**
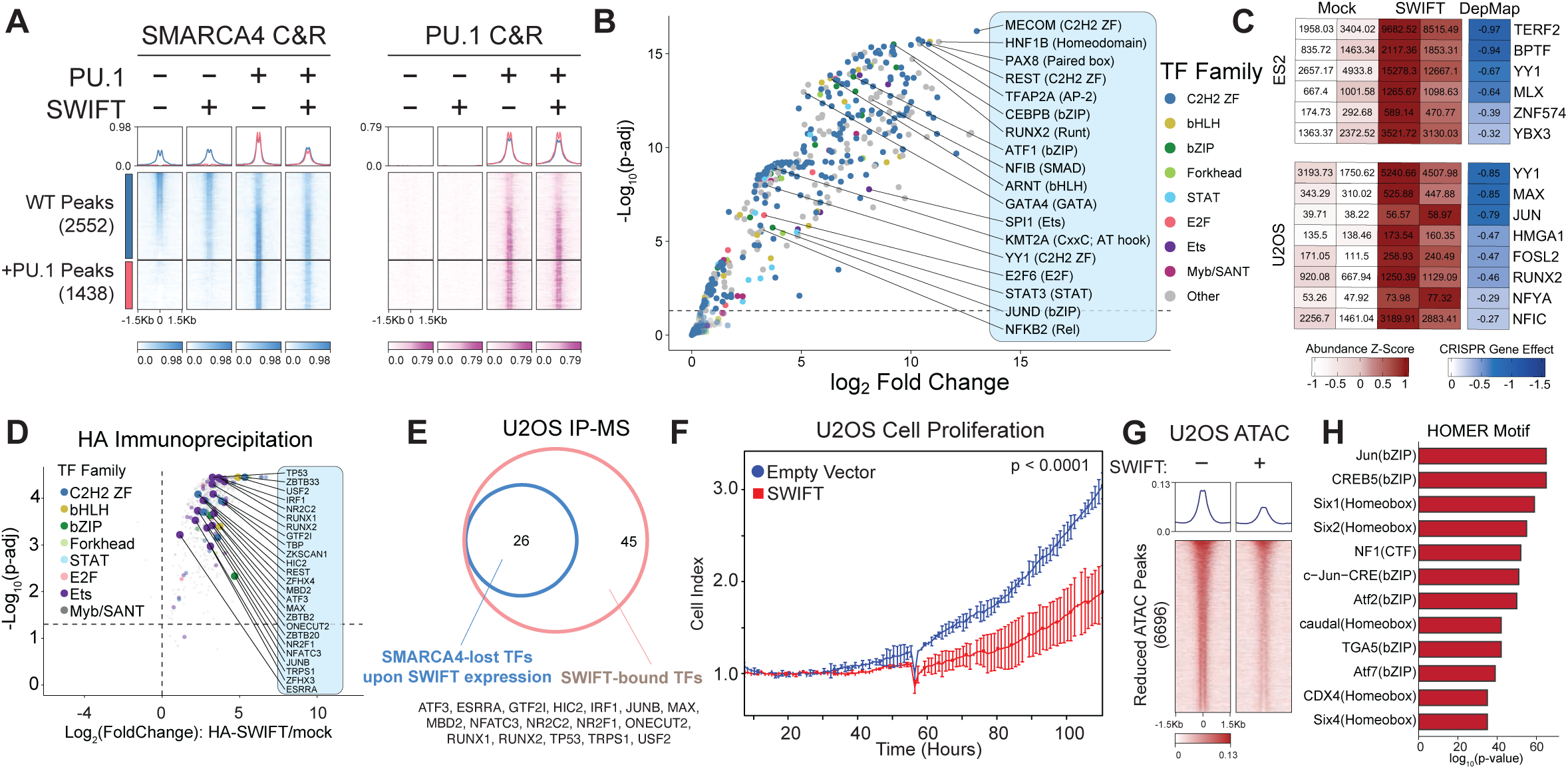
The SWIFT domain is an evolutionarily conserved SWI/SNF-specific platform for diverse transcription factor interactions. **(A)** Heatmap displaying the CUT&RUN-determined occupancy of SMARCA4 and PU.1 in hMSCs expressing PU.1 in the presence or absence of SMARCD2 SWIFT domain expression. **(B)** Scatterplot displaying the composite enrichment of transcription factors bound by 200 nM SMARCD2 SWIFT (versus mock control) incubated in nuclear extracts isolated from U2OS osteosarcoma, ES2 and OVISE ovarian carcinoma, and NCI-H1048 small cell lung cancer cell lines as detected by mass spectrometry (n= 2 replicates for each cell line). **(C)** Heatmaps displaying the enrichment of TFs bound by SMARCD2 SWIFT from ES2 and U2OS cell nuclear extracts (mock shown as control) and their CRISPR gene dependency scores (DepMap). **(D)** Scatter plots showing enrichment of TFs, colored by family, in SWIFT domain pulldown (left) in the setting of HA-SWIFT overexpression in U2OS cells. TFs that also showed reduced binding to mSWI/SNF in the same cells are shown in larger circles. **(E)** Venn diagram depicting overlap between TFs enriched (gained) in SMARCD2 SWIFT domain IP-MS and those reduced in mSWI/SNF (SMARCA4) binding. Selected TFs are labeled. **(F)** Proliferation of U2OS cells overexpressing SMARCD2 SWIFT domains, monitored by eSight. P-value p<0.0001 of difference in the slope of exponential growth for each line (n= 3 replicates) is indicated. **(G)** Heatmap showing the ATAC-Seq signal at 6,696 sites of reduced accessibility in U2OS cells expressing HA-tagged SMARCD2 SWIFT domain. **(H)** Homer motif enrichment at sites indicated in G.

Evolutionary analyses of the primary structure of SWIFT domain and structural conservation shows that it is a SWI/SNF-specific Ig-fold, evolved in the earliest SWI/SNF complexes in yeast and maintained throughout phylogeny (**Fig. S9C**). This YEATS-like domain is substantially distinct from the YEATS domains in other chromatin modifiers (AF9, GAS41, etc.) (**fig. S5C-E**) (*81*, *83*, *84*). Notably, in the yeast SMARCD homolog, SNF12, the YEATS domain-specific aromatic residues that are necessary for interaction with histone H3 acylation are also missing, suggesting distinct evolution of SWIFT and YEATS domains (*85*).

The evolutionary conservation of SWIFT led us to hypothesize that it functions as a broad TF-binding platform beyond PU.1. To this end, we incubated purified HA-tagged SMARCD2 SWIFT with nuclear extracts from five distinct cancer cell lines, followed by HA-immunoprecipitation and mass spectrometry (MS) (**Fig. 5B, and fig. S9D-E)**. Intriguingly, we detected significant enrichment of several TFs that co-immunoprecipitated with the SMARCD2 SWIFT domain including YY1, JUN, FOSL2, and RUNX2, relative to mock control (**Fig. 5C, and fig. S9E**). Intriguingly, incubation of the OCA-T2 TF showed binding with SMARCD2 SWIFT domain, but did not co-immunoprecipitate with the YEATS domains of AF9 and GAS41, supporting functional divergence (**fig. S9F-G).** Importantly, R290W mutation or treatment of the SWIFT domain with a small molecule inhibitor of its hydrophobic pocket, PPSC1 (*99*), reduced binding to majority of the TFs (**fig. S9H-J**), suggesting that many TFs interact with the SWIFT domain through the same hydrophobic pocket as PU.1. These results are consistent with the shared chemical properties of TF TADs that lack conservation of primary structures.

To identify the SWIFT domain binding site on the ubiquitous AP.1 family TF, FOS, we assessed a series of FOS deletion mutants (**fig. S10A-B**). We found that three separate 80-aa deletions on the C-terminal tAD of FOS reduced SWIFT engagement (**fig. S10B**). Additionally, three 50-aa long fragments showed independent binding to the SWIFT domain, with a Leucine/Serine-rich C-terminal region showing the strongest binding (**fig. S10A, C**). These results extend our findings with PU.1 that tADs can interact with SWIFT through allovalent interactions.

Notably, several SWIFT domain-interacting TFs represented top dependencies in cancer cell lines in which interactions were identified, such as BPTF and TERF2 in ES cells and YY1 and MAX in U2OS sarcoma cells (**Fig. 5C**). Further, we expressed SWIFT in isolation in U2OS cells and performed IP-MS studies to identify factors enriched on HA-SWIFT and simultaneously decreased in abundance on mSWI/SNF complexes via SMARCA4 IP. Notably, TFs such as RUNX1, RUNX2, TP53, among others, engaged with the dominantly expressed HA-SMARCD2 SWIFT domain and showed reduced engagement with endogenous mSWI/SNF complexes, absent changes in mSWI/SNF abundance (**Fig. 5D-E and fig. S10D**).

With this, we hypothesized that ectopic expression of the SMARCD SWIFT domain in isolation could “poison” the required binding platform for TFs upholding cancer cell proliferation by blocking the interaction of essential TFs with the SWIFT domain of endogenous mSWI/SNF complexes. Indeed, overexpression of the SMARCD2 SWIFT domain in U2OS osteosarcoma and ES2 ovarian cancer cell lines significantly attenuated proliferation (**Fig. 5F, and fig. S10E**). Consistent with an essential role of the hydrophobic pocket for interaction with TFs, expression of R290W mutant SMARCD2 SWIFT domain did not impair cell proliferation relative to WT SWIFT (**Fig. S10F-H)**). Proliferative attenuation via SWIFT domain overexpression was comparable to mSWI/SNF SMARCA4/2 ATPase inhibition using FHD-286 (**fig. S10I**). Mechanistically, we identified 6,696 sites at which genomic accessibility (ATAC-Seq signal) was reduced upon overexpression of SMARCD2 SWIFT (**Fig. 5G**). Indeed, these sites contained motifs of TFs which are dependencies in U2OS cells (such as JUN) and have previously been shown to be necessary for their proliferation (**Fig. 5H**)(*100*). Together, these data highlight that exogenous expression of SWIFT dominantly blocks endogenous mSWI/SNF complexes from interacting with TFs, resulting in failure of TF-mediated mSWI/SNF activity and oncogenic proliferation.

### TFs display SMARCD paralog-specific preference for interactions with SWIFT domains

Mammalian cells have three highly homologous SMARCD paralogs – SMARCD1, D2, and D3 – that incorporate into mSWI/SNF complexes in a mutually exclusive manner (**Fig. 6A, and fig. S11A**). The SMARCD1 and SMARCD2 paralogs are generally ubiquitously expressed in most human tissues, with variation by tissue type, while SMARCD3 expression is lower and more restricted to a select set of cell types such as cardiomyocytes, cerebellum, ovarian tissues and monocytes (**fig. S11B-C**).

**Figure 6.**
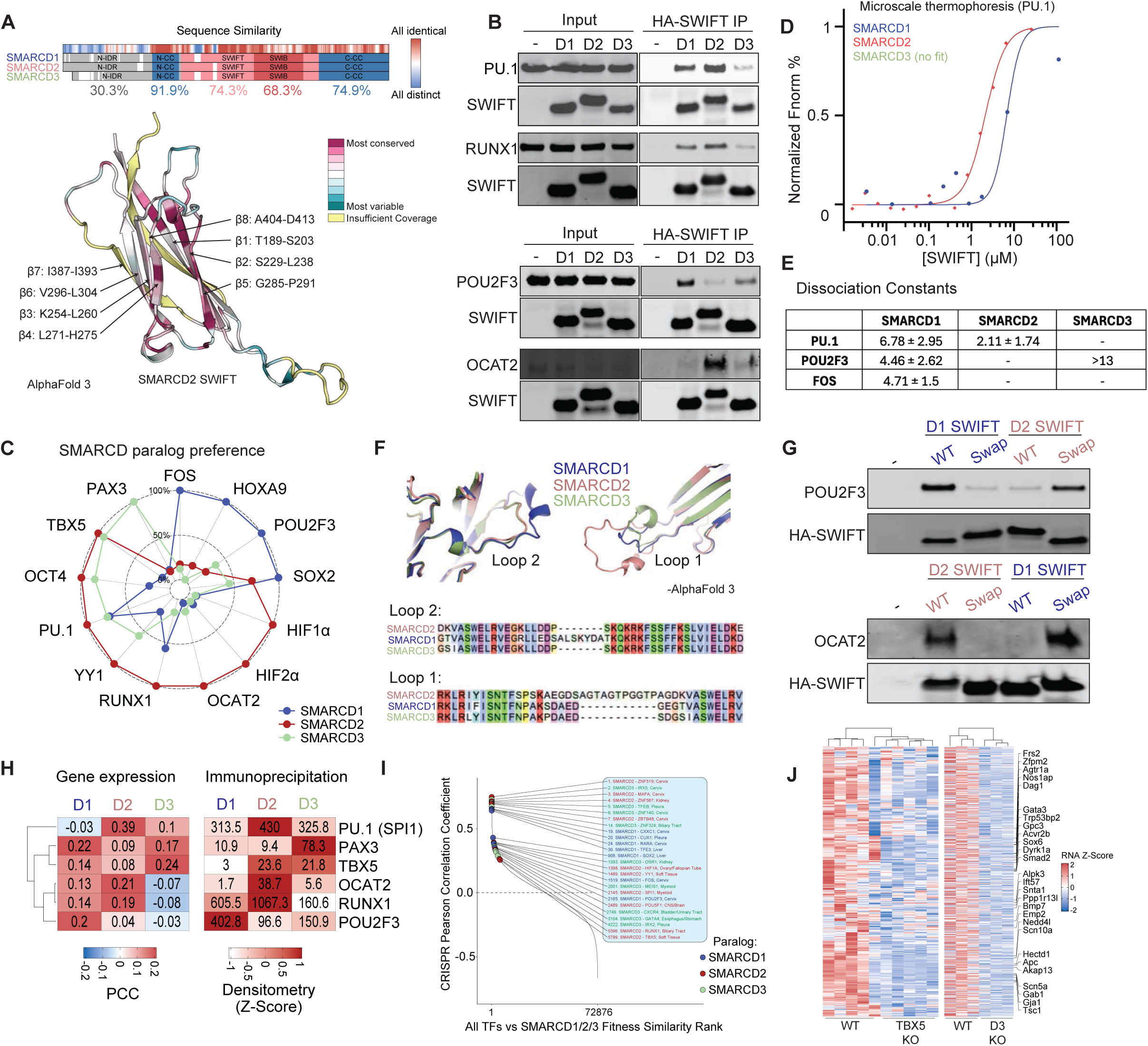
Transcription factors preferentially interact with paralog-specific SMARCD SWIFT domains. **(A)** Domain structure and sequence conservation of human SMARCD paralogs (top). Evolutionary sequence conservation score of SMARCD SWIFT domain is mapped on its structure in divergent colors (bottom). **(B)** Immunoblots of HA-SWIFT immunoprecipitation from HEK293T cells co-expressing HA-tagged SWIFT of specific SMARCD paralog (D1, D2, D3) and the indicated FLAG-tagged transcription factors. **(C)** Circle plot showing the preferential interaction between the tested transcription factors and paralog-specific SWIFT domains. Fluorescence intensities of TF/SWIFT were normalized across SMARCD paralogs and are plotted as a fraction of maximum intensity. **(D)** SWIFT was titrated into 5 nM of PU.1 for microscale thermophoresis to measure dissociation constants of interactions between paralog-specific SWIFT domains and PU.1. **(E)** Table showing the dissociation constants of PU.1, POU2F3 and FOS for SWIFT domains of SMARCD1/2/3 paralogs. NA: fit not found. **(F)** Overlap of predicted structures of SMARCD2 and SMARCD3 SWIFT domains (salmon and green, respectively) on cryoEM structure of SMARCD1 SWIFT (blue), showing the structural divergence of loop 2 and loop 1 within the paralogs. Linear sequence alignment is shown below. **(G)** Immunoblots of eluates from HA-SWIFT immunoprecipitation from HEK293T cells expressing POU2F3 (top) or OCAT2(bottom) and SMARCD1/D2 SWIFT domains containing wildtype loops or all of their loops swapped. **(H)** Heatmaps showing the Pearson Correlation Coefficients of gene expression patterns of SMARCD paralogs and lineage-specific TFs across all cell lines (left). Paralog-specificity as determined by z-score normalized intensities of TF immunoblot signal from HA-SWIFT immunoprecipitation are shown on right. **(I)** Rank plot showing all TF-SMARCD gene pairwise fitness correlations from DepMap CRISPR studies performed across ∼1000 cancer cell lines. Selected top-ranked SMARCD1/2/3-TF correlations are labeled. **(J)** Heatmap showing the gene expression profiles of cardiomyocyte-specific genes regulated by TBX5 transcription factor in WT cardiomyocytes and TBX5 knockout cells (left), compared to WT cardiomyocytes and SMARCD3 KO cells (right).

To probe the differences in interactions between various TFs and SMARCD paralogs, we ectopically co-expressed them at equivalent levels in HEK-293T cells and performed HA-SMARCD1/2/3 immunoprecipitation. Surprisingly, despite the high sequence and structural conservation between the SWIFT domains of the three paralogs, TFs displayed variation in their paralog-specific binding preferences (**Fig. 6B-C, and fig. S11D**). Similarly, incubation of the SWIFT domain of SMARCD1/2/3 paralogs with ES2 nuclear extracts enriched distinct TFs to varying degrees (**fig. S12A-B**). For example, RUNX1 and PU.1 showed preference for the SMARCD2 SWIFT domain and to a lesser extent that from SMARCD1, while PAX3 and TBX5 showed preference for the SMARCD3 SWIFT (**Fig. 6B-C**). The majority of the TFs evaluated co-immunoprecipitated with SMARCD2 or SMARCD1 subunits, which are the most broadly expressed in human tissues (**fig. S11B-C**). While 5 of 14 TFs bound SMARCD3 to some degree, only PAX3 displayed a strong SMARCD3-specific preference. Notably, inclusion of the SWIB domain on loop 7 (aa312-382) did not alter SMARCD paralog specific preference of TFs such as OCA-T2, suggesting that the preference is encoded within the SWIFT domain (**fig. S12C-D**). Finally, we determined dissociation constants for select TF-SWIFT domain interactions. Indeed, PU.1 that showed preference for the SMARCD2 SWIFT exhibited the lowest KD for the SMARCD2 SWIFT interaction (KD: 2.11 ± 1.74 µM for SMARCD2, 6.78 ± 2.95 µM for SMARCD1, no fit for SMARCD3) whereas POU2F3 strongly prefers the purified SMARCD1 SWIFT domain (KD: 4.46 ± 2.62 µM for SMARCD1, >13 µM for SMARCD3) (**Fig. 6D-E**). Mirroring the binding preference observed in cells, FOS only showed interaction with SMARCD1 at KD of 4.71 ± 1.5 µM (**fig. S12E**).

The three SMARCD paralogs are highly homologous in the structured β-sheets that form the β-sandwich. However, the L1 and L2 loops, which are the longest flexible loops in the SWIFT domain between the first three N-terminal β-strands, are the most diverse between homologs (**Fig. 6F, and fig. S11A)**. In particular, SMARCD1 has longer L2 loops, which make a unique, short alpha helix. SMARCD2 and SMARCD3 paralogs lack these residues and instead have Proline242 that prevents formation of a secondary structure. In contrast, SMARCD2 contains a longer L1 loop, absent in SMARCD1 and SMARCD3, which imparts a higher net positive surface charge to it near the hydrophobic pocket.

To evaluate whether flexible loops in SWIFT impart the paralog-specific preferences for interactions with TFs, we created SMARCD1 and SMARCD2 mutants in which we swapped the flexible loops between the paralogs and tested their interaction with POU2F3 and OCAT2 TFs, the two TFs that displayed the strong preferences for either SMARCD1 or SMARCD2, respectively. Indeed, we found that the SMARCD1 SWIFT containing SMARCD2 loops immunoprecipitated markedly reduced levels of POU2F3 (**Fig. 6G**). Reciprocally, swapping in SMARCD1 loops was sufficient to impart complete binding of POU2F3 to SMARCD2 SWIFT (**Fig. 6G**). Similarly, the SMARCD2 SWIFT containing SMARCD1 loops completely lost ability to bind OCAT2, while SMARCD2 loops swapped into SMARCD1 SWIFT domain enabled the OCAT2-SMARCD1 SWIFT interaction. Together, these data indicate that unstructured loops between the β-strands confer subunit paralog-specific binding of TFs.

Our data support a model in which specific SMARCD paralogs are required to accommodate specialized lineage-specific TFs to control site-specific targeting in cancer cell contexts. To directly test this hypothesis, we expressed HA-tagged SMARCD1 or SMARCD2 in U2OS cells and probed occupancy of SMARCA4 at de novo sites (**fig. S12F-G**). Motif enrichment analyses under SMARCD1- and SMARCD2-specific loci revealed several unique TF motifs for each paralog (**fig. S12H**). Consistent with our biochemical results (**fig. S12H**), motifs for RUNT (RUNX2) and ETS family TFs (ERS) were enriched in SMARCD2-specific peaks.

Finally, the surprising binding preferences of TFs for specific SMARCD subunit paralogs of SMARCD SWIFT domains prompted us to evaluate a potential molecular role enabled by these biochemical interactions. We noted that expression of tissue-specific TFs, such as PU.1, RUNX1, PAX3 and TBX5 correlates strongly with the expression of their biochemically preferred SMARCD-homolog across human tissues (**Fig. 6H, and fig. S12I**). Further, we evaluated all 72,876 pairwise fitness similarity correlations derived from DepMap across ∼1000 cancer cell lines between human TFs and SMARCD1, D2, or D3 paralogs and identified several paralog-specific TF-SMARCD correlations (**Fig. 6I**).

In addition, we integrated two RNA-seq datasets from cell culture-based models of cardiomyocyte differentiation (*101*, *102*) to test the model that SMARCD paralog switches enable affinities for specific TFs to control cell state-specific mSWI/SNF targeting required for proper cellular differentiation. Indeed, *SMARCD3* expression is activated specifically during cardiomyocyte differentiation, along with several cardiomyocyte-specific TFs including *TBX5* and *GATA4* **(fig. S12J)**. SMARCD3 knockout cells fail to turn on the cardiomyocyte-specific gene expression signature, regulated by TBX5 (**Fig. 6J, and fig. S12K**), suggesting that SMARCD3-containing mSWI/SNF complexes facilitate TBX5-target gene expression. Our result that TBX5 preferentially interacts with SMARCD3 SWIFT provides a biochemical explanation for the previously observed molecular cooperativity between TBX5 and SMARCD3 during cardiomyocyte differentiation (*24*, *103*, *104*).

Taken together, these results indicate that switches in the subunit composition of mSWI/SNF complexes, a longstanding and well appreciated phenomenon during development (*17*, *20*, *24*, *36*, *104–109*), at least in part, facilitate this critical biochemical mechanism to regulate TF-driven gene regulatory trajectories hallmark to cell differentiation or oncogenesis, thus presenting a mechanism by which combinatorial diversity encoded by the mSWI/SNF complexes is essential for transcriptional regulation.

## Discussion

The mSWI/SNF core module, which lacks sequence-specific DNA-binding domains and histone reader domains, is the first to assemble (*11*), suggesting that direct interaction-based mechanisms beyond these modes guide complex localization across cells and tissue types, particularly to specialized distal enhancer sites(*110*). Previous studies have explored interactions between TFs and mSWI/SNF complexes through immunoprecipitation and genomic co-localization studies. Here, we discover a platform on mSWI/SNF complexes, the SWIFT domain, that directly interacts with TFs encompassing diverse families, including those considered pioneer, settler, lineage-specific or ubiquitous.

Using PU.1-mSWI/SNF as an example, we show that multiple binding sites within a tAD interact with the same binding pocket on SWIFT. Such allovalent protein-protein interactions have been well characterized for interactions of protein kinases, phosphorylation readers and DNA-binding domains of TFs (*111–116*), where multiple binding sites on a protein trap binding partners within its IDR (*111*, *112*, *114*, *115*, *117–119*). Akin to this mode of interaction, the SWIFT domain is a binding platform for a wide range of TFs, which, while lacking a conserved linear sequence within TADs, have shared “grammar” of amino acid composition– disordered domains with biased representation of hydrophobic, acidic and Q-rich amino acids. During the preparation of our manuscript, a preprint by Sadalge et. al. also characterized an interaction between a PU.1 peptide (aa 60-100) and the SMARCD subunit of mSWI/SNF complexes at the region we identify and term SWIFT via its hydrophobic pocket, consistent with our study (*149*). However, they propose a ‘lock-and-key’ binding mechanism between a single site in PU.1 that binds to the SWIFT domain through specific interactions. Such specific interactions between PU.1 residues and SWIFT requires a highly specific primary structure within the tADs of TFs and is inconsistent with our finding that SWIFT interacts with many diverse TFs. Therefore, our results suggest that the PU.1-BAF interaction characterized in Sadalge et. al. is indeed one specific configuration out of many that TFs can adopt to interact with SWIFT through their divergent tADs; biochemical studies presented here identify a broad range of TF-SWIFT interactions.

Previous studies have identified that switches in the expression of SMARCD paralogs facilitate appropriate differentiation trajectories (*24*, *102*, *103*, *106*, *107*). Surprisingly, we discovered that TFs display a high degree of preference SMARCD paralog-specific SWIFT domains and that this specificity is conferred by divergent loops within the SWIFT domain (**Fig. 6F-G**). It remains unclear what sequence features within the TF tADs underlie their preference for specific SMARCD paralogs. Future studies are needed to comprehensively determine in vitro affinities across TFs with each SMARCD paralog SWIFT domain. Nonetheless, we find that TF-SMARCD paralog SWIFT binding preferences for select lineage-specific TFs mirrors their co-expression across cell types (**Fig. 6H**). Our results provide a biochemical basis for the model that mSWI/SNF compositions switch during development, here suggesting SMARCD paralog switches fine-tune complex affinities for lineage-specific TFs to facilitate divergent cellular differentiation trajectories.

Finally, CRISPR screens have revealed that several cancers exhibit dependencies on specific SMARCD paralogs. For example, SMARCD3 supports oncogenic transcription in a subset of medulloblastomas (Group 3, ∼30% of all medulloblastoma), pancreatic adenocarcinoma, and gastric cancers(*108*, *109*, *120*). Our data suggest that TF-SWIFT interactions and hence TF-mediated mSWI/SNF recruitment underlie these results. Further biochemical and structural characterization of specific TF-SWIFT interactions may inform therapeutic opportunities to manage TF-driven cancers with enhanced precision relative to broad mSWI/SNF inhibition such as via FHD-286, a clinical-grade dual SMARCA2/4 ATPase inhibitor (*121*, *122*). Our findings here open new possibilities for targeted inhibition of TF-addicted cancers via SWIFT domain disruption.

## Supporting information

Data S1. GEE Labeling

Data S2. AlphaFold Error values

Data S3. nHDX-MS

Data S4. Immunoprecipitation-MS

Data S5. SWIFT pulldown TMT-MS

Data S6. SWIFT paralog pulldown TMT-MS

## Acknowledgements

We thank members of the Kadoch Laboratory for productive discussions and assistance throughout the duration of this project. We also thank Zach Herbert and Maura Sullivan of the DFCI Molecular Biology Core Facility (MBCF) for help with high-throughput sequencing and the Harvard Medical School Center for Macromolecular Interactions (CMI) for assistance with MST experiments. We thank Dr. Clayton Collings for bioinformatic support and implementation of new analytical approaches. We also thank the following sources for support for this work: National Institutes of Health New Innovator Award 1DP2CA195762 (C.K.); Pew-Stewart Scholars in Cancer Research Award (C.K.); American Cancer Society Research Scholar Award RSG-14-051-01-DMC (C.K.); The Mark Foundation for Cancer Research Emerging Leader Award (C.K.); The Gabrielle’s Angel Foundation Award (C.K); The Howard Hughes Medical Institute (C.K.); National Institutes of Health Grants 5R01CA259365 (C.K.); National Institutes of Health Grant R01AG079283 (M.L.G.); National Institutes of Health Grant R24GM136766 (M.L.G); National Institutes of Health Grant K99CA237855 (N.M.); National Institutes of Health Grant R01GM132129 (J.A.P.); National Institutes of Health Grant R35 GM156406 (J.A.P.); National Institutes of Health Grant GM067945 (S.P.G.); AACR John and Elizabeth Family Foundation Basic Science Fellowship (S.U.J.); National Science Foundation Graduate Research Fellowship Program (K.E.W.)

## Author Contributions

Conceptualization: S.U.J., K.E.W., C.K.; Methodology: S.U.J, K.E.W., A.W.Y., K.S., A.M.T., J.A.P., N.M., C.K.; Investigation: R.J.J., K.S., S.R., M.P.A., A.M.T., D.S.G., J.A.P., N.M., H.W.R., C.F.L.; Data Analysis, Statistics, Visualization: A.W.Y. and A.S.; Funding acquisition: C.K.; Supervision: M.P. M.L.G., C.K.; Writing – original draft: S.U.J., K.E.W., C.K.; Writing – review & editing: S.U.J., K.E.W., A.W.Y., C.K.

## Data and materials availability: Data and materials availability

All sequencing data are made available in the Gene Expression Omnibus (GEO) database under the series GSE310119; http://www.ncbi.nlm.nih.gov/geo/GSE310119. All raw proteomics and mass-spectrometry datasets have been deposited to ProteomeXchange via PRIDE database under accession number PXD070375. Processed mass-spectrometry data are available as supplementary files. For HDX, original mass spectra and the protein sequence database used for searches have been deposited in the public proteomics repository MassIVE (http://massive.ucsd.edu) and are accessible at ftp://MSV000099556@massive-ftp.ucsd.edu. Processed mass spectrometry data is available as supplementary files. No new code was generated for this work; all analyses have been performed according to the details described in the Methods section using available software packages. Reagents generated in this study are available from the corresponding author upon reasonable request and completion of a material transfer agreement.

## Materials and Methods

### Cell lines and cell culture

HEK293T lentiX (Clonetech) cells were grown in Dulbecco’s modified Eagle’s medium (Gibco) supplemented with 1% GlutaMax (Gibco), 1% sodium pyruvate (Gibco), 1% mouse embryonic fibroblast non-essential amino acids (Gibco)), 1% Penicillin/Streptomycin (Gibco) with 10% fetal bovine serum (Gibco). Cells were passaged every 3-4 days as described above. Human ASC52telo, hTERT immortalized adipose derived mesenchymal stem cells (MSCs) (ATCC) were grown in mesenchymal stem cell basal medium for adipose, umbilical and bone marrow derived MSCs (ATCC) supplemented with mesenchymal stem cell growth kit for adipose and umbilical-derived MSCs - Low Serum (ATCC) at 37°C, 5% CO2, 95% humidity according to the manufacturer’s protocol. Cells were split every 3-4 days and resuspended with fresh medium to maintain culture confluency between 60-80%. To passage, cells were washed with pre-warmed PBS pH 7.4 (Gibco Life Technologies 10010-049) and detached using room temperature TrypLE Express Enzyme (Life Technologies). Cells were passaged with fresh media 20X before a new batch of cells were used. BIN67 cells (gift of Barbara Vanderhyden) were grown in custom media (40% Dulbecco’s modified Eagle’s medium, 40% Ham’s F12 (Gibco), 20% fetal bovine serum (FBS) (Gibco) supplemented with 1% Glutamax (Gibco) and 1% Penicillin-Streptomycin (Gibco)) at 37°C, 5% CO2, 95% humidity. MOLM-13 and Ovise cells were grown in RPMI-1640 (Gibco), 10% fetal bovine serum (Gibco), supplemented with 1x Glutamax (Gibo), 1% Penicillin-Streptomycin. ES2 and U2OS cells were cultured in EMEM media (Gibco) with 10% fetal bovine serum (Gibco), supplemented with 1x Glutamax (Gibo), 1% Penicillin-Streptomycin. Cells were maintained by passaging every 4-5 days when cells reached 80-90% confluency by washing with pre-warmed PBS pH 7.4 and detaching cells with Trypsin-EDTA (0.25%) (Gibco).

### Cloning of Mammalian and Bacterial Expression Constructs

All TF expression constructs were PCR-amplified from cDNA or existing expression constructs purchased from Addgene, Dharmacon, or obtained from Harvard University DNA repository and the DNASU Plasmid repository using Phusion High-Fidelity DNA Polymerase (NEB) with GC buffer (NEB) or HF buffer (NEB) and custom PCR primers introducing an in-frame N-terminal HA-tag (YPYDVPDYA). PCR-amplified fragments were gel purified using 1% agarose gel in Tris-Acetate EDTA (TAE) buffer and PCR cleanup columns (Qiagen) and cloned into a modified pTight vector from Clonetech (EFL1-alpha promoter) with puromycin resistance or blasticidin resistance cassette using In-fusion (Clonetech) at the Not1 cut site. Recombination products were transformed into One-Shot Stbl3 chemically competent E. coli (Invitrogen) and colonies were selected for plasmid purification (Qiagen) and sequence verified. BRG1 WT-V5 and BRG1 K785R-V5 lentiviral expression constructs in pLEX307 were constructed as previously published(*33*, *68*). EF-1a-MCS-PGK- HA-DPF2 construct used for endogenous cBAF purification was previously described(*9*, *11*). PU.1 deletion mutant constructs were constructed by PCR amplifying fragments flanking the desired deletion with 30 bp of complementarity. N-terminal and C-terminal fragments were amplified separately, gel purified, and fused together using a second PCR with the two fragments as templates. Final fragments were gel purified and cloned into a modified pTight vector (EF1-alpha promoter) containing blasticidin resistance, as above. Single amino acid alanine substitution mutations were generated by site directed mutagenesis (SDM) using Q5 Site-Directed Mutagenesis kit (NEB) according to manufacturer’s recommendations using primers designed by NEBaseChanger (NEB). Constructs for bacterial expression and purification of mutants were cloned into pGOOD with N-terminal GST and C-terminal 6X His tags using In-fusion (Clontech) at the EcoRI cut site. pMD2.G was a gift from Didier Trono (Addgene plasmid # 12259 ; http://n2t.net/addgene:12259 ; RRID:Addgene_12259). psPAX2 was a gift from Didier Trono (Addgene plasmid # 12260 ; http://n2t.net/addgene:12260 ; RRID:Addgene_12260). Human SWIFT domain sequence was obtained from IDT and cloned into pGOOD with N-terminal 6x-His tag for protein purification from bacterial cells. For protein purification from mammalian cells, SWIFT domain sequence with N-terminal FLAG tag and C-terminal HA-tag was cloned into pCAG plasmid (addgene 11150) using infusion master mix (Clontech).

### Protein sequences in this study

#### SMARCD1 SWIFT

IKQKRKLRIFISNTFNPAKSDAEDGEGTVASWELRVEGRLLEDSALSKYDATKQKRKFSSFFKSLVIELDKDLYGPDNHLVEWHRTATTQETDGFQVKRPGDVNVRCTVLLMLDYQPPQFMPPEPIIINHVISVDPNDQKKTACYDIDVEVDDTL

#### SMARCD2 SWIFT

LTQKRKLRIYISNTFSPSKAEGDSAGTAGTPGGTPAGDKVASWELRVEGKLLDDPSKQKRKFSSFFKSLVIELDKELYGPDNHLVEWHRMPTTQETDGFQVKRPGDLNVKCTLLLMLDHQPPQYQHPDPIVINHVISVDPNDQKKTACYDIDVEVD

#### SMARCD3 SWIFT

MKQKRKLRLYISNTFNPAKPDAEDSDGSIASWELRVEGKLLDDPSKQKRKFSSFFKSLVIELDKDLYGPDNHLVEWHRTPTTQETDGFQVKRPGDLSVRCTLLLMLDYQPPQFLPPDPIVINHVISVDPSDQKKTACYDIDVEVEEPL

### Lentiviral Production and Transduction

Lentiviral particles were produced by transfecting HEK293T LentiX (Clontech) cells plated to 80% confluency using 1 µg mL^-1^ PEI (Polysciences, Inc) and lentiviral expression vector and packing vectors psPAX2 and pMD2.g in ratio of 4:3:1 respectively as previously described(*123*, *124*). 72 hours after transfection, media was filtered through 0.45 µm filter (EMD Millipore SE1M003M00) and viral particles were concentrated by ultracentrifugation at 20,000 rpm for 2.5 hrs at 4°C using a SW32Ti rotor. Viral particles were resuspended in PBS and used to transduce cells plated to 70% confluency using 8 µg ml^-1^ polybrene (Santa Cruz Biotechnology) or 10 µg mL^-1^ protamine sulfate (Sigma-Aldrich, P4020-5G) for BIN-67. Media was exchanged 24 hours following virus addition and cells were selected with 8 µg mL^-1^ blasticidin or 1 µg mL^-1^ puromycin starting 48 hours after virus addition. Expression of desired constructs was verified by western blot and cells were harvested for genomics based analyses on Day 7 post-transduction.

### Nuclear Extraction and Immunoprecipitation

Nuclear protein extracts were prepared using standard techniques. Briefly, cells were washed with cold PBS to remove media. Pelleted cells were resuspended in hypotonic buffer (EBO: 50mM Tris pH7.5, 0.1% N-40, 1 mM EDTA, 1 mM MgCl2, supplemented with protease inhibitors (10 µg mL^-1^ each chymostatin, pepstatin, leupeptin (EMD Millipore) in DMSO and 1mM PMSF (Gold Bio) in isopropanol). Nuclei were pelleted at 5,000 rpm for 5 min at 4°C. Nuclei were resuspended in high salt buffer (EB300: 50mM Tris pH 7.5, 300mM NaCl, 1% NP-40, supplemented with protease inhibitors as above). Lysates were incubated on ice for 10 min and sonicated using 3-6 pulses of 10 seconds with a Misonix Sonicator 3000 Ultrasonic Cell Disruptor system with microtip probe with max power output of 6W. Lysates were pelleted for 5 min at max speed in a benchtop centrifuge and supernatant was quantified using BCA assay reagents (Pierce). 0.5-1 mg of protein per condition were supplemented with 1 mM DTT (Sigma-Aldrich) and was used for immunoprecipitation with 3-5 ug of antibodies overnight at 4°C with rotation. To monitor transient interactions between TFs and BAF complexes, lysate was diluted with low salt buffer (EB0, 50 mM Tris pH 7.5, 0 mM NaCl, 1% NP-40) to final 150 mM NaCl. 30 µl of Protein-G Dynabeads were added and incubated for 2 hours with rotation and washed with EB150 six times and eluted with 2X LDS loading dye with 1 mM DTT at 95°C for 5 min and loaded on SDS-PAGE gel. Whole cell lysates were prepared using 5X pellet volume of protein extraction buffer (20mM Tris, 1.5% SDS). Lysed cells were incubated at 95°C for 2 min to denature DNA, sonicated and immunoblotted. Immunoprecipitations were repeated 2-3 times.

### SWIFT domain immunoprecipitation for probing paralog-specific preferences for TFs

Plasmids containing genes for FLAG-tagged transcription factors and HA tagged D1, D2, D3 SWIFT domains were co-transfected into Expi293F cells (thermo scientific; Cat# A14527) grown in Expi media (thermo scientific; Cat# A1435102). 20ug of plasmids (pCAG containing gene of interest) (10ug of TF and 10ug of SWIFT) were transfected into 30mL of culture using 20ug of polyethyleminine. After 16 hours, valproic acid, sodium propionate, 1x NEAA, and 1x GlutaMAX were added to culture. Cells were harvested after 2 days by centrifugation at 1000 g for 10 min, washed with PBS and resuspended in lysis buffer (20 mM HEPES pH 8.0, 1 mM EDTA, 200 mM KCl, 0.1% NP-40, 1x protease inhibitor cocktail, 0.5 mM DTT, 0.4 mM PMSF). After protein extraction for 1 hour and clarification using centrifugation at 20,000 g for 30 min, whole cell extract was diluted 1:1 with HEK0 buffer (20 mM HEPES pH 8.0, 1 mM EDTA, 1x protease inhibitor cocktail, 0.5 mM DTT, 0.4 mM PMSF). Lysates were incubated with 30ul of Pierce™ Anti-HA Magnetic Beads (Thermo Scientific) for 2.5 hours. Beads were washed 4 times with HEK150 buffer (20 mM HEPES pH 8.0, 1 mM EDTA, 150 mM KCl, 0.05% NP-40, 1x protease inhibitor cocktail, 0.5 mM DTT, 0.4 mM PMSF), 2 times with lysis buffer (20 mM HEPES pH 8.0, 1 mM EDTA, 200 mM KCl, 0.1% NP-40, 1x protease inhibitor cocktail, 0.5 mM DTT, 0.4 mM PMSF), and once with PBS. Beads were eluted in 100 ul 2X SDS for western blotting. Most immunoprecipitations were performed once; some key experiments such as loop swap experiments were performed three times.

### SDS-PAGE and western blotting

Western blotting was performed using standard techniques. Briefly samples were loaded on 4-12% Bis-Tris Plus SDS-PAGE gels (Invitrogen) and run at 80V for 20 min, followed by 130V for 1 hour. Proteins were transferred to polyvinylidene difluoride (PVDF) membranes (Immobilon-FL, EMD Millipore) for 90 min at 30V and blocked for 1 hr at room temperature in 5% milk (Boston BioProducts) in PBS. Blots were incubated O/N with primary antibodies, washed 3X in PBST and incubated with secondary fluorophore-conjugated species-specific antibodies (Li-Cor) for one hour. Membranes were washed 3X with PBS-T and 1X with PBS and visualized using Li-Cor Odyssey CLx.

### CUT&RUN

Concanavalin A (ConA) beads (Polysciences, Cat#: 86057-10) were activated with Bead Activation Buffer containing 20 mM HEPES pH 7.9, 10 mM KCl, 1 mM CaCl2, 1 mM MnCl2; beads were stored on ice until used. 500,000 cells per sample were harvested and resuspended in media at 1 million cells per mL. Cells were lightly crosslinked with 16% paraformaldehyde at a final concentration of 0.1% at room temperature for 2 minutes. Crosslinking was quenched with 2.5M Glycine at a final concentration of 200mM at room temperature for 5 minutes. Cells were pelleted by centrifugation (300 rcf, 5 minutes, room temperature) and were washed with PBS containing 200mM Glycine. Cells were resuspended with 100ul/sample of Nuclear Extraction Buffer containing 20 mM HEPES pH 7.9, 10 mM KCl, 0.1% Triton X-100, 20% Glycerol supplemented with fresh 0.5 mM Spermidine and 1X protease inhibitor (Sigma Aldrich, Cat# 11836170001) for 2 min on ice. Nuclei were pelleted by centrifugation (600 rcf, 3 minutes, 4C), and resuspended in 100ul/sample cold Wash150 Buffer containing 20 mM HEPES pH 7.9, 150 mM NaCl, 1% Triton X-100, 0.05% SDS supplemented with fresh 0.5 mM Spermidine and 1X protease inhibitor (Sigma Aldrich, Cat# 11836170001) and incubated with activated ConA beads at room temperature for 15 minutes. Nuclei-ConA bead complexes were resuspended in 50 ul/sample cold Antibody Buffer containing 20 mM HEPES pH 7.9, 150 mM NaCl, 1% Triton X-100, 0.05% SDS, 2mM EDTA supplemented with fresh

0.5 mM Spermidine, 1X protease inhibitor (Sigma Aldrich, Cat# 11836170001), 0.01% Digitonin, and 0.5 ug primary antibody/sample, and incubated overnight nutating at 4C. Following removal of supernatant, ConA-nuclei complexes were washed twice with Digitonin Buffer containing 20 mM HEPES pH 7.9, 150 mM NaCl, 1% Triton X-100, 0.05% SDS supplemented with fresh 0.5 mM Spermidine, 1X protease inhibitor (Sigma Aldrich, Cat# 11836170001), and 0.01% Digitonin. Samples were then resuspended in 50ul Digitonin Buffer and 2 ul of CUTANA pAG-MNase (Epicypher, Cat#: 15-1116) each and incubated at room temperature for 10 minutes. Samples were washed twice with Digitonin Buffer and resuspended in 50ul Digitonin Buffer and 1ul of 100mM CaCl2 per sample. Following incubation nutating at 4C for 2 hours, samples received 33ul of Stop Buffer containing 340 mM NaCl, 20 mM EDTA, 4 mM EGTA supplemented with fresh 50 ug/mL RNase A, 50 ug/ml Glycogen and incubated at 37C for 1 hour to release MNase digested DNA fragments. Supernatant was collected into fresh tubes and supplemented with 0.8uL 10% SDS and 1uL of 20mg/mL Proteinase K for overnight incubation at 55C. DNA was purified with Ampure XP (Beckman Coulter, Cat#: A63881) beads and 80% EtOH washes. DNA was eluted with 25uL of 0.1X TE buffer and transferred to new tubes. DNA fragments were prepared for next generation sequencing with CUTANA CUT&RUN Library Prep Kit (Epicypher, Cat#: 14-1001) according to manufacturer protocol. CUT&RUN experiments were performed in one replicate but with antibodies for different mSWI/SNF subunits. Overlap between distinct subunit peaks were used for high confidence in results.

### ATAC-seq Sample Preparation

ATAC-seq was performed as previously described(*125*) with 100,000 cells per sample. Cells were dissociated and counted, as described above, 7-days post transduction with lentivirus. Cells were washed with cold PBS, resuspend in lysis buffer (10mM Tris-HCl pH7.4, 10 mM NaCl, 3 mM MgCl2, 0.1% IGEPAL) and centrifuged at 500 g for 10 min at 4°C. Nuclei were resuspended in transposition mix (1X TD Buffer, 2.5 ul TDE1 Nextera Tn5 Transposase (Illumina)) and incubated with mild rotation for 30 min at 37°C. Tagmented DNA was purified using MinElute PCR cleanup kit (Qiagen). Samples were amplified using 10 total PCR amplification cycles with custom PCR Primers using NEBNext HF PCR Master Mix (NEB). Samples were purified with 1.2X Agencourt AMPure XP beads (Beckman Coulter) and size distribution was determined using D5000 HS reagents and screentapes (Agilent) on 2200 TapeStation Instrument (Agilent). Samples were pooled in equimolar amounts and sequenced on Illumina Next-seq 500 with NextSeq 500/550 High Output Kit V2 75 cycles (Illumina) using 37bp paired-end sequencing parameters. ATAC-Seq experiments were performed in two replicates.

### RNA-seq Sample Preparation

Cells were dissociated and counted, as described above, 7-days post transduction with lentivirus. 2 × 10^6^ live cells as were washed with PBS and RNA was extracted using QIAshredder and RNeasy Mini Kit (Qiagen) per the manufacturer’s instructions. 1 µg of total RNA was used as input for library preparation and processed per the manufacturer’s instructions using NEBNext Poly(A) mRNA magnetic isolation module (NEB) followed by the NEBNext Ultra II Directional RNA library prep kit for Illumina (NEB) with NEBNext Multiplex Oligos for Illumina (NEB). Samples were collected in biological replicate. ERCC synthetic RNA spike-in control (Thermo Fisher Scientific) was added at the beginning of library preparation per manufacturer’s recommendations to normalize samples across conditions. Libraries were quantified using Qubit dsDNA HS Assay Kit (Thermo Fisher Scientific) and size distribution was verified using Tapestation D1000 HS reagents and screentapes (Agilent). Equimolar amounts of libraries were pooled, denatured, and sequenced on Illumina Next-seq 500 with NextSeq 500/550 High Output Kit V2 75 cycles (Illumina) according to standard Illumina protocols with single-end sequencing parameters. RNA-Seq were performed in two biological replicates.

### CUT&TAG

CUT&TAG was performed following published protocols (Epicypher)(*126*). Cells were dissociated from culture plates using TrypLE Express (Thermo Fisher Scientific) and washed once with PBS pH 7.4 (Gibco). Cells were counted with Countess II (Invitrogen). 200,000-500,000 live cells were used per epitope as determined by Trypan Blue (Thermo Fisher Scientific) exclusion. Cells were processed for Cut & Tag as previously published(*127*) and according to manufacturer’s CUTANA protocol (Epicypher) with the use of BioMag Plus Concanavalin A beads (Polysciences), CUTANA pAG-Tn5 (Epicypher), primary antibodies, and Guinea Pig anti-Rabbit IgG (Heavy & Light chain) antibody (Antibodies-Online). Tn5 transposed fragments were amplified between 14-18 cycles using NEB Next HF 2X PCR Master Mix (NEB) and custom primers originally designed for ATAC-seq library amplification to allow for multiplexing of 50+ samples. Amplified DNA was purified using 0.75X AMPpure Beads (Beckman Coulter) to remove primer-dimers. Libraries were quantified and QC’d as above. Samples were sequenced by the Dana Farber Cancer Institute Molecular Biology Core Facility using paired-end read parameters on an Illumina Nextseq 500 (Illumina) using Illumina NextSeq 500/550 High Output Kit V2 75 cycles (Illumina).

### cBAF Purification from Mammalian Cells

Purification of endogenous mammalian cBAF complexes using HA-DPF2 as bait and purified HA-PU.1 was performed as previously described (*11*) using hypotonic followed by high salt lysis to extract nuclear proteins. Briefly, stable cell lines expressing lentivirally transduced HA-tagged baits were cultured in 150mm dishes and expanded to 300 (HA-DPF2) or 150 (HA-PU.1) total plates. For purifications of cBAF containing SMARCD2 WT or R290W, cells expressing N-terminal V5-tagged SMARCD2 constructs were used. Cells were scraped and washed with cold PBS. Cells were lysed in hypotonic buffer (HB: 10 mM Tris HCl pH 7.5, 10 mM KCl, 1.5 mM MgCl2 1 mM DTT, and 1 mM PMSF) for 5 min on ice and centrifuged at 5000 rpm for 5 min at 4°C. Pellets were resuspended in HB buffer supplemented with protease inhibitor cocktail and cells were lysed using glass Dounce homogenizer. Lysed cells were centrifuged at 5000 rpm for 20 min at 4°C and nuclei were resuspend in High Salt Buffer (HSB: 50mM Tris HCl pH 7.5, 300 mM KCl, 1mM MgCl2, 1 mM EDTA, 1% NP40, 1mM DTT, 1 mM PMSF and protease inhibitor cocktail) and incubated with rotation for 1 hour at 4°C followed by centrifugation to remove chromatin pellet at 20,000 rpm for 1 hour at 4°C using SW32Ti rotor. Chromatin pellet was removed and supernatant was filtered through Advantec Grade QR200 Quartz Fiber Filters (Cole-Parmer). Complexes were purified from clarified nuclear extracts using magnetic HA beads (Pierce) overnight at 4°C. HA beads were washed 6X with HSB and purified proteins were eluted using Elution Buffer (HSB + 1 mg/ml HA peptide (GenScript) 4X for 1.5 hours each at 4°C. Eluted proteins were subjected to silver staining and density gradient sedimentation (see below) for analysis of yield and purity.

### Density sedimentation gradients

Eluted protein complexes were separated by density using 10%–30% glycerol gradients prepared using Gradient Master containing 25mM HEPES pH 7.9, 0.1mM EDTA, 12.5mM MgCl2, 100mM KCl supplemented with1mM DTT and protease inhibitors as previously described (*9*, *11*). Briefly, eluted proteins were loaded on top of 11 ml gradient and centrifuged at 40,000 rpm for 16 hours at 4°C. Individual fractions (21 × 550 µl) were manually aspirated from the top of the gradient. 80 µL of each collected fraction were concentrated using 10 µL of Strataclean beads (Agilent), eluted with 2X NuPage LDS sample buffer (ThermoFisher Scientific), loaded onto SDS-PAGE gels and stained using Silver Quest Silver Staining Kit (ThermoFisher Scientific) according to manufacturer’s protocol. Fractions with confirmed purified expression of desired protein/complex were pooled and concentrated using appropriate molecular weight cutoff protein concentrator (Pierce). Samples were snap frozen and kept at -80°C until needed.

### Protein Expression and purification of TFs

Transcription factors (PU.1, FOS, and POU2F3) for in vitro binding experiments was purified from Expi293F cells (thermo scientific; Cat# A14527), grown in Expi media (thermo scientific; Cat# A1435102). 800 µg of plasmids (pCAG containing gene of interest) were transfected into 2 billion cells in 1 L culture using 4 mg of polyethyleminine. After 16 hours, valproic acid, sodium propionate, 1x non-essential amino acids were added to culture. Cells were harvested after 2 days by centrifugation at 1000 g for 10 min, washed with 50 ml of PBS and resuspended in lysis buffer (20 mM HEPES pH 8.0, 1 mM EDTA, 300 mM KCl, 1% NP-40, 1x protease inhibitor cocktail, 0.5 mM DTT, 0.4 mM PMSF). After protein extraction for 1 hour and clarification using centrifugation at 20,000 g for 30 min, whole cell extract was incubated with 0.7 ml of FLAG M2 magnetic beads (Sigma; Cat# M8823) for 2 hours. Beads bound to proteins of interest were washed 6-times with binding buffer and proteins were eluted using 3x FLAG peptides.

Eluted proteins were further purified by anion exchange chromatography using a custom Capto-Q column (0.15 ml column volume) and eluted by a KCl (0-1 M) over 25 column volumes. Fractions containing proteins of interest were pooled and further purified by size-exclusion chromatography using superdex 200 (Cytiva; Cat# 28-9909-44). Fractions containing protein of interest were pooled and used for downstream experiments.

### Protein Disorder Prediction

Disordered region prediction in Fig. 2A was performed using metapredict (*128*, *129*), accessed using the web portal (metapredict.net), using the canonical sequence for PU.1 (Uniprot: P17947). Disordered regions were identified as having a disorder score greater than 0.5, and was plot using ggplot2.

### Microscale Thermophoresis

Transcription purified from Expi293T cells using FLAG affinity purification followed by captoQ anion exchange chromatography and superdex 200 size-exclusion chromatography were labeled with lysine-reactive 2^nd^ generation Red NHS labeling kit (Nanotemper, Cat# MO-L011). 5 nM labelled TFs were incubated with varying concentration of SMARCD1/2/3 SWIFT domains in binding buffer containing 20 mM HEPES pH 8.0, 100 mM KCl, 0.5 mM EDTA, 1 mM DTT, 0.05% tween-20, and 0.1 mg/ml BSA for 1 hour at 4 °C and 30 min at room temperature. Samples were filled into premium capillaries (Nanotemper, Cat# MO-K025) and microscale thermophoresis was performed on Monolith NT.115 (C-303). Raw data was imported into Graphpad Prism 10 and Hill Curve was fitted into data to identify EC50 values (analogous to concentration of SWIFT at which half the TFs are bound, equivalent to KD for a one-site binding). For data visualization of paralog-specific fits, Fnorm values were normalized to the bottom and top saturation values obtained from the Hill curve. Each experiment was performed in two independent replicates; data is shown for a representative curve.

### Peptide Pulldowns

Peptide pulldowns were performed as previously described (*130*). 3 nmol of biotinylated PU.1 peptides (novopep, custom) were incubated with 0.3 nmol of purified HA-tagged SMARCD2 SWIFT or AF9 YEATS domains overnight in binding buffer containing 20 mM HEPES, 1 mM EDTA, 75 mM KCl, 0.01% tween-20. 25 µl of streptavidin conjugated Dynabeads (cat# 65601) for 2 hours to capture biotinylated peptides. Dynabeads were washed 7-times with binding buffer and boiled in 2x Laemmli buffer to elute captured proteins. After removal of Dynabeads, eluates were analyzed by immunoblotting using HA antibodies (Cell Signaling; Cat# 3724). Peptide pulldowns were performed once and were repeated for some key peptides with SWIFT containing WT and R290W with similar results.

### Nuclear extract preparation and SWIFT-pulldown experiments

For SWIFT pulldown experiments, 1-2 billion cells were resuspended in hypotonic lysis buffer (20 mM HEPES pH 8.0, 1 mM EDTA, 4 mM MgCl2, 10 mM KCl, 1x protease inhibitor cocktail, 1 mM DTT, 0.4 mM PMSF). Nuclei were resuspended in 10 ml of buffer-AB (20 mM HEPES pH 8.0, 110 mM KCl, 1 mM EDTA, 2 mM MgCl2, 1x protease inhibitor cocktail, 0.4 mM PMSF, 1 mM DTT). 1.1 ml of 4 M ammonium sulfate was added to nuclei suspension. After 1 hour incubation, chromatin fraction was pelleted by ultracentrifugation in Ti-45 rotor at 35k RPM for 90 min. Nuclear proteins were salted out by slowly adding 0.3 g of powdered ammonium sulfate to the clarified extract followed by ultracentrifugation at 20K RPM in Ti-45 rotor for 30 min. Pellet containing nuclear proteins were resuspended in binding buffer containing 20 mM HEPES pH 8.0, 150 mM KCl, 1 mM EDTA, 0.5 mM DTT, 1x protease inhibitor cocktails, 0.4 mM PMSF. Purified N-terminally HA-tagged SWIFT domain was incubated with nuclear extract at a final concentration of 200 nM for 2 hours at 4 °C. Following binding, HA-antibody conjugated dynabeads (Thermo scientific; Cat# 88837), pre-blocked with BSA were added to the mixture for 2 hours at 4 °C. Beads were washed 6 times in binding buffer and stored in 200 mM HEPES pH 8.0 until mass-spectrometry. Experiments were performed in two independent replicates.

### Electrophoretic Mobility Shift Assay

The protein-DNA binding reaction was carried out as previously published with a few modifications (*131*). Short DNA probes containing PU.1 DNA binding motif or scrambled control were ordered as custom single stranded oligos from Integrated DNA Technologies (IDT). Equimolar amounts of forward and reverse oligos were mixed in annealing buffer (20 mM HEPES pH 7.5, 50 mM NaCl, 1 mM EDTA), denatured at 95 °C for 5 min and annealed by slowly cooling to 16 °C (Table S3). 200 nM of purified HA-PU.1 was incubated with 100 nM of IRdye800 labeled PU.1-binding oligo and 500 nM unlabeled scrambled probes in 10 µl reaction buffer containing 50 mM HEPES pH 7.5, 50 mM KCl, 1 mM MgCl2, 1mM EDTA, 5% glycerol. cBAF was titrated into the reaction and incubated at room temperature for 30 min after which reaction was mixed with 10X DNA loading dye and run on 8% Novex TBE gel (ThermoFisher Scientific) in 1X TBE buffer at 100V for 1 hour followed by 30 min staining with 1X SybrGold in TBE and visualization using DNA station. These experiments were performed two times with similar results.

### GEE labeling

Differential GEE footprinting was performed on purified cBAF complexes, cBAF complexes incubated with nucleosome core particle (NCP), and cBAF complexes incubated with purified PU.1. The molar ratios for cBAF complexes with nucleosome and PU.1 were 1: 5 and 1: 8, respectively. To reach the binding equilibrium, cBAF complexes and its binding partners were incubated under 4°C overnight, prior to the footprinting. For GEE labeling, glycine ethyl ester (GEE), 1-ethyl-3-(3-dimethylaminopropyl) carbodiimide (EDC) stock solutions were prepared fresh in PBS buffer. GEE was added to each pre-equilibrated samples, followed by addition of EDC which initiates the footprinting reaction. The final concentrations of protein, GEE, and EDC were 250 nM, 20 mM, 1 mM, respectively. The reaction was carried out at 25°C for 1 h, before quenching by addition of equal volume of 1 M ammonium acetate. The proteins were then immediately purified by acetone precipitation. Labeling experiments were performed in triplicate n=3.

### Proteolysis of GEE labeled peptides

Urea (8M) was added to dissolve the acetone-precipitated protein pellets. The proteins’ cysteine residues were then reduced and alkylated with TCEP and iodoacetamide (IAM), respectively. Lys-C and trypsin were next successively added with a enzyme-to-protein ratio of 1:20 and 1:5 (w:w), respectively. Samples were incubated at 37 °C with Lys-C for 6 h and trypsin for 12 h. The digestion was quenched by adding formic acid to a final concentration of 5% (by volume).

### Mass spectrometry analysis of GEE labeled peptides

A Dionex UltiMate 1000 system (Thermo Fisher Scientific) was coupled to an Orbitrap Fusion Lumos (Thermo Fisher Scientific) through an EASY-Spray ion source (Thermo Fisher Scientific). Nanoflow liquid chromatography separation was carried as previously described(*95*). Briefly, peptide samples were loaded (15 µl min^−1^; 1 min) onto a trap column (100 µm × 2 cm; 5 µm Acclaim PepMap 100 C18; 50 °C), eluted (0.2 µl min^−1^) onto an EASY-Spray PepMap RSLC C18 column (2 µm; 50 cm × 75 µm ID; 50 °C; Thermo Fisher Scientific) and separated with the following gradient (all % buffer B (that is, 0.1% formic acid in acetonitrile)): 0–110 min: 2–22%; 110–120 min: 22–35%; 120–130 min: 35–95%; 130–150 min: isocratic at 95%; 151–153 min: 95–2%; 153–171 min: isocratic at 2%. The spray voltage was 1,900 V, the ion transfer tube temperature was 275 °C and the RF lens was 30%. Mass spectrometry scans were acquired in profile mode and tandem mass spectrometry scans were acquired in centroid mode, for ions with charge states 2–7, with a cycle time of 3 s. Mass spectrometry spectra were recorded from 375–1,500 Da at 120-K resolution (at *m/z* 200), and higher-energy collisional dissociation tandem mass spectrometry was triggered above a threshold of 2.0 × 10^4^, with quadrupole isolation (0.7 Da) at 30-K resolution and a collision energy of 30%. Dynamic exclusion was used (60 s), and monoisotopic precursor selection was on.

### GEE labeling mass -spectrometry data processing

Identification of the unmodified peptides and assigned modifications were done by using Byologic (Protein Metrics) and further validated by manual inspection. Modification sites were identified based on MS/MS. Signal intensities of the unmodified peptide (Iu) and its modified species (Iox) were integrated using Byologic (Protein Metrics) from the extracted ion chromatograms (XICs). The GEE modification fraction of a residue was calculated using the following equation: % modified = Imodified/(Imodified + Iunmodified)×100. Quantification of the modified species was based on the GEE products and its hydrolyzed products (+85.0522 Da, +57.0209 Da

### Mapping changes in GEE exposure onto 3D mSWI/SNF structures

Changes in exposure between the bound state (cBAF+NCP or BAF+PU.1) and unbound state (cBAF only) were visualized on published cryoEM structure cBAF complex. Change values were extended in windows of 11aa centered on each GEE label and values were averaged in overlapping windows. Ambiguous GEE labels (labels with more than one candidate D/E site) were split into multiple observations and preprocessed similarly. GEE labels from paralogs not represented on the structure (ACTA2, ARID1B, BCL7B, BCL7C, SMARCD2, SMARCD3) were remapped to the primary paralog in the structure (ACTA2:ACTB, ARID1B:ARID1A, BCL7B:BCL7A:, BCL7C:BCL7A, SMARCD2:SMARCD1, SMARCD3: SMARCD1), and preprocessed and visualized similarly. Changes in exposure were stored in the b-factor column of the PDB file and visualized using the third-party color_b PyMOL script as a blue-white-red heatmap clipped at -30% (dark blue) to 30% (bright red).

### Mapping changes in GEE exposure to cBAF complex subunit protein schematics

Changes in exposure between the bound state (cBAF+NCP or cBAF+PU.1) and unbound state (cBAF only) were preprocessed as described in 3D Structure. However, paralogs were not remapped to the primary paralog resolved in the structure of the endogenous cBAF complex to show paralog-specific effects (ARID1A vs ARID1B, SMARCD1 vs SMARCD2, SMARCC1 vs SMARCC2). Change values were visualized as vertical lines across the length of each BAF subunit (blue as a loss in exposure and red as a gain in exposure upon substrate binding). Changes in exposure in cBAF-NCP and cBAF-PU.1 data for the cBAF-NCP structure were merged into the same plot using matplotlib.

### cBAF-NCP and cBAF-GEE Overlap Calculations

Changes in exposure visualized across the lengths of the cBAF subunits were analyzed by eye to define merged sites (windows that were within 10-20aa of each other with consistent effects were merged) of exposure. Merged windows of exposure change less than 1% were omitted as noise. Overlapping merged sites of exposure with consistent effects (i.e. both gains in exposure or loss of exposure) when bound to either substrate, NCP and PU.1, were considered overlapped sites of exposure. The remaining sites were binned into disjoint sets of cBAF+NCP exclusive sites and cBAF+PU.1 exclusive sites. These counts were visualized as a Venn diagram using matplotlib.

### Distribution of Exposure between the ATPase and Core Modules

Changes in exposure were filtered to the most significant sites using a 5% cutoff for cBAF+NCP and cBAF+PU.1. Gains and losses in exposure within the Core and ATPase modules for both GEE datasets were tallied and reported as proportions using stacked bar charts in matplotlib.

### Hydrogen-Deuterium Exchange (HDX)

HDX-MS experiments were conducted on the fully automated Chronect HDX extended parallel platform (Trajan Scientific and Medical) coupled to an NCS-3500RS Nano system and an Orbitrap Exploris™ 480 mass spectrometer (Thermo Fisher Scientific). Stock solutions of PU.1 were prepared in 20 mM HEPES (pH 7.0), 150 mM NaCl, 0.5 mM EDTA, and 5 mM TCEP. For deuterium labeling experiments, PU.1 was diluted to a final concentration of 2.2 µM in the absence or presence of 11 µM D1_SWIFT. The exchange reaction was initiated by diluting 3 μL of the protein sample with 9 μL of a D₂O-based buffer (20 mM HEPES pH 7.0, 150 mM NaCl, 0.5 mM EDTA, and 5 mM TCEP), resulting in a final D₂O concentration of 67%. Samples were incubated at 4 °C for three distinct time points: 20, 100, and 1000 seconds. The exchange reactions were terminated by the addition of 50 μL of ice-cold quench buffer (4 M Guanidine hydrochloride, 0.5% Formic Acid, pH 2.3) at 0 °C. A fraction of this sample (18 μL) was injected for subsequent online digestion using a dual-protease column containing pepsin and protease XIII (1:1, Novabioassays) for 3 min at 200 μL/min. Peptides were separated using a 12 minute gradient and analyzed in the MS via a data-independent acquisition (DIA) workflow as described previously(*132*). Deuterium incorporation was quantified using HDExaminer™ PRO (version 0.9.0.104, Trajan Scientific and Medical). Samples were run in quadruplicates. Significant differences in % deuterium uptake at the peptide level were identified using a dual-threshold approach: (1) a two-tailed Welch’s t-test at α = 0.05, and (2) a difference threshold of 2 times the pooled standard deviation from % deuterium uptake measurements across peptides that do not pass the Welch’s t-test significant cutoff.

### RNA-seq Data Analysis

RNA-Seq reads were demultiplexed using bcl2fastq v2.20.0.422 aligned to the hg19 genome with STAR v2.5.2b(*133*). Upregulated and downregulated genes were determined using DESeq2 (log2FC = 1, B-H p-value = 0.05)(*134*). DESeq2’s estimateSizeFactors function was used to generate normalized counts (*135*). ggplot2 was used to generate volcano plots for changes in expression visualized as scatter plots. The eulerr R package was used to generate venn diagrams of differential genes. ComplexHeatmap R package was used to generate heatmaps and visualize Z-Score normalized read counts for each gene across all conditions for each cell line. The Kendall and Spearman distance functions were used to perform hierarchical clustering in heatmaps. clusterProfiler GSEA function (*136*) was used through the msigdbr R package to perform GSEA analysis using the Hallmark and C2 gene sets. The “stat” output from DESeq2 was used to determine Differential expression ranking. MOLM13 sgPU.1 RNA data was downloaded from GEO series GSE186131(*137*). For fig. S2D, the Benjamini-Hochberg method was applied to adjust p-values and adjusted p-values are reported.

For Fig. 6G, RNA-seq expression data from cardiomyocytes (day 10), left atria tissue, and left ventricle tissue were obtained from GSE59383 (*102*), GSE129503 (*138*), and GSE222579 (*101*), respectively. Provided raw gene counts were renormalized using DESeq2’s DESeqDataSetFromMatrix, and Z-score values were calculated by dataset for visualization purposes. The top tertile of differential genes by Z-score were calculated for each dataset and used to plot Z-scores for Tbx5 KO and Smarcd3 KO, where each row represents a gene concordantly changing in all three datasets.

### ATAC-seq Data Analysis

ATAC-seq reads were first trimmed using fastp v0.24.0, aligned with Bowtie2 v2.2.9 to the GRCh38 reference genome, and filtered using SAMtools v0.1.19 to remove duplicates and for quality (-F 256 -f 2 - q 30) (*139–141*). Reads mapping to regions defined in the ENCODE project’s unified GRCh38 blacklist bed file were removed using bedtools v2.30.0[14]. ATAC-seq data was processed by merging technical replicates using SAMtools merge. Peaks were called using MACS3 (-f BAMPE -g hs -q 0.001 --nomodel --extsize 200 –broad)(*142*).

Differential accessibility was determined by calculating reads-per-million values across a merged set of peaks for a given comparison set, and using an RPM fold-change cutoff of 1.5. Differential accessibility was then visualized with venn diagrams generated using eulerr. Site centers with flanking windows of 200bp (total window size of 400bp) were identified across given FASTA sequence sites. Motif enrichment across these sets of sites was determined using HOMER findMotifsGenome.pl against genone-background (-size 400) unless otherwise noted. Barplots were generated to visualize HOMER motif results using ggplot2. Heatmaps and deepTools computeMatrix as used to generate metaplots over indicated peaks(*143*). deepTools bamCoverage (--binSize “40” --normalizeUsing “CPM” --exactScaling) was used to generate bigwig inputs for heatmaps. Stacked bar charts generated with ggplot in R to highlight distance to TSS highlighting the proportion of promotor, promoter proximal and distal enhancer regions, determined using bedtools and the GENCODE v46 annotations (*144*).

### CUT&RUN and CUT&TAG Data Analysis

ChIP-seq reads were aligned in a manner identical to ATAC-seq reads. Peaks for CUT&RUN were generated with MACS2 (-f BAM -g hs -q 0.01 --nomodel). Peaks for CUT&TAG were generated using SEACR (at a q value of 0.01, using the “non” normalized and “stringent” settings)(*145*).

FASTA sequences across these sets of sites were generated using site centers with flanking windows of 200bp (total window size of 400bp). Enriched motifs across these sets of sites were determined using HOMER findMotifsGenome.pl against genome-background (-size 400) unless otherwise noted. Bigwigs were generated using deepTools and rendered using karyoplotR.

### AP-MS Analysis

Differential proteins from AP-MS data were identified using DeqMS and plotted via ComplexHeatmap and ggplot2. Differential transcription factors were identified using the list of transcription factors published in Lambert et al(*58*).

### GTEX and CCLE Analysis

Analysis of CCLE and DepMap CRISPR screening data was conducted using the public 25q2 release(*43*, *146*). Analysis of GTEX expression data was conducted using version 10.

### Cryo-EM Structures and Alphafold Predicted Structures

Resolved Cryo-EM structures for 9A0K, 6LTJ, 4TMP, 3DO7, and 1FRG were downloaded from PDB and rendered using Pymol(*9*, *71–73*, *80*). Electrostatic surfaces were also generated using Pymol. Alphafold2 predictions for the proteome were used to determine structural domains and were also used to add unresolved domains to 6LTJ via alignment(*72*).

Alphafold3 predictions were used to predict affinity between the SMARCD2 SWIFT domain and PU.1 sequences(*147–150*). Predictions were parsed to extract the contact probability between each residue within a subsequence and L256, I258, V272, W274, F286, and V288 of SMARCD2, and aggregated as a log2 mean probability, which was then averaged across all sequence predictions containing that residue. Alphafold3 was accessed at alphafoldserver.com

### Sequence and Structure Alignment

Sequence alignment was conducted using Clustalw (*151*) using the primary isoform of the SwissProt entry for a given protein. Sequence conservation was determined using the sequence of the isolated SMARCD2 SWIFT domain and Consurf. Structural similarity was determined using Foldseek (*152*) to search for the Alphafold3 predicted structure of the SMARCD2 SWIFT domain against the Uniprot/Proteome database. Protein hits identified in this manner were then aligned against each other in an all-to-all fashion to obtain a TMscore, which was then used to perform hierarchical clustering and displayed using dendextend(*153*).

