## Supplementary Material for "A SWI/SNF-specific Ig-like domain, SWIFT, is a transcription factor binding platform"

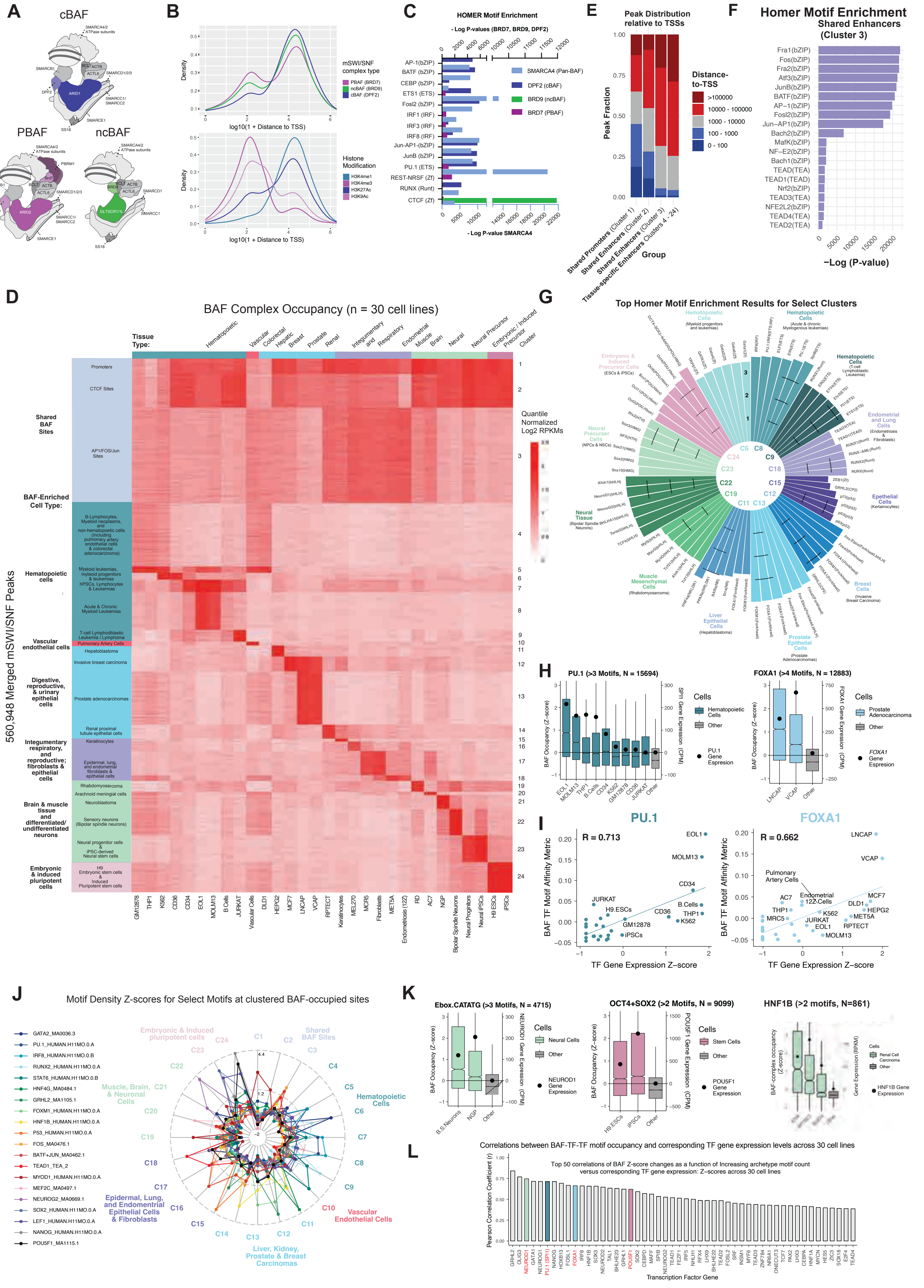

A
cBAF
PBAF
ncBAF
B
mSWI/SNF complex type
PBAF (BRD7)
ncBAF (BRD9)
cBAF (DPF2)
Histone Modification
H3K4me1
H3K4me3
H3K27Ac
H3K9Ac
Density
log10(1 + Distance to TSS)
C
HOMER Motif Enrichment
- Log P-values (BRD7, BRD9, DPF2)
SMARCA4 (Pan-BAF)
DPF2 (cBAF)
BRD9 (ncBAF)
BRD7 (PBAF)
- Log P-value SMARCA4
E
Peak Distribution relative to TSSs
Peak Fraction
Distance-to-TSS
>100000
10000 - 100000
1000 - 10000
100 - 1000
0 - 100
Group
Shared Promoters (Cluster 1)
Shared Enhancers (Cluster 2)
Shared Enhancers (Cluster 3)
Tissue-specific Enhancers Clusters 4 - 24)
F
Homer Motif Enrichment
Shared Enhancers
(Cluster 3)
−Log (P-value)
D
BAF Complex Occupancy (n = 30 cell lines)
Tissue Type:
Cluster
Quantile Normalized Log2 RPKMs
560,948 Merged mSWI/SNF Peaks
Shared BAF Sites
BAF-Enriched Cell Type:
Hematopoietic cells
Vascular endothelial cells
Digestive, reproductive, & urinary epithelial cells
Integumentary respiratory, and reproductive; fibroblasts & epithelial cells
Brain & muscle tissue and differentiated/ undifferentiated neurons
Embryonic & induced pluripotent cells
G
Top Homer Motif Enrichment Results for Select Clusters
H
PU.1 (>3 Motifs, N = 15694)
FOXA1 (>4 Motifs, N = 12883)
BAF Occupancy (Z-score)
SPI1 Gene Expression (CPM)
FOXA1 Gene Expression (CPM)
Cells
Hematopoietic Cells
Other
PU.1 Gene Espression
Prostate Adenocarcinoma
FOXA1 Gene Espresion
I
PU.1
FOXA1
R = 0.713
R = 0.662
BAF TF Motif Affinity Metric
TF Gene Expression Z−score
J
Motif Density Z-scores for Select Motifs at clustered BAF-occupied sites
GATA2_MA0036.3
PU.1_HUMAN.H11MO.0.A
IRF8_HUMAN.H11MO.0.B
RUNX2_HUMAN.H11MO.0.A
STAT6_HUMAN.H11MO.0.B
HNF4G_MA0484.1
GRHL2_MA1105.1
FOXM1_HUMAN.H11MO.0.A
HNF1B_HUMAN.H11MO.0.A
P53_HUMAN.H11MO.0.A
FOS_MA0476.1
BATF+JUN_MA0462.1
TEAD1_TEA_2
MYOD1_HUMAN.H11MO.0.A
MEF2C_MA0497.1
NEUROG2_MA0669.1
SOX2_HUMAN.H11MO.0.A
LEF1_HUMAN.H11MO.0.A
NANOG_HUMAN.H11MO.0.A
POU5F1_MA1115.1
Embryonic & Induced pluripotent cells
Shared BAF Sites
Hematopoietic Cells
Muscle, Brain, & Neuronal Cells
Epidermal, Lung, and Endometrial Epithelial Cells & Fibroblasts
Vascular Endothelial Cells
Liver, Kidney, Prostate & Breast Carcinomas
K
Ebox.CATATG (>3 Motifs, N = 4715)
OCT4+SOX2 (>2 Motifs, N = 9099)
HNF1B (>2 motifs, N=861)
Neural Cells
Stem Cells
Other
NEUROD1 Gene Expression
POU5F1 Gene Expression
HNF1B Gene Expression
L
Correlations between BAF-TF-TF motif occupancy and corresponding TF gene expression levels across 30 cell lines
Top 50 correlations of BAF Z-score changes as a function of Increasing archetype motif count versus corresponding TF gene expression: Z-scores across 30 cell lines
Pearson Correlation Coefficient (r)
Transcription Factor Gene

**Fig. S1**: **Unique sequence-specific transcription factor motifs underlie cell-type specific mSWI/SNF complex occupancy at distal enhancers genome-wide. A.** Schematic depicting the subunit compositions of cBAF, PBAF and ncBAF complexes. **B.** Density plots showing the distribution of cBAF, PBAF and ncBAF peaks (top) and histone modifications (bottom panel) as a function of distance from annotated transcription start sites (x-axis). **C**. Homer motif enrichment -Log (p-value) for top 15 enriched transcription factor motifs over sites occupied by SMARCA4 (pan-BAF), DPF2 (cBAF), BRD7 (PBAF) and BRD9 (ncBAF) by ChIP-seq in the EOL1 cell line. **D**. Unsupervised clustering of 560,948 merged BAF complex-occupied ChIP-seq peaks (SMARCA4/SMARCA2) genome-wide across 30 human cell lines/cell types. Cell lines are annotated by tissue and cancer types. **E**. Distance from TSS of mSWI/SNF peaks in clusters defined in D. **F.** Homer motif enrichment -Log (p value) for top enriched transcription factor motifs over sites in cluster 3 as defined in D. **G.** Homer motif enrichment (log (p-value)) of top transcription factor motifs in tissue-specific mSWI/SNF clusters defined in D. **H**. BAF complex occupancy (Z-score) at sites with >3, >4 TF motifs for PU.1 and FOXA1, respectively, in cell lines of shared lineage where corresponding TF gene expression is >1 CPM. Expression of the PU.1 and FOXA1 in the cell lines is marked in black circle. **I.** Correlation between BAF occupancy as a function of corresponding TF motif archetype counts (Z-score (SMARCA4 ChIP-seq RPKM / [Motif])) for motifs of PU.1 and FOXA1, respectively in cell lines of shared lineages and their expression (z-score normalized). **J.** Motif density Z-scores for transcription factors at various clusters of BAF complex occupancy. **K.** BAF complex occupancy (Z-score) at sites with >3, >2, and >2 TF motifs for NEUROD1, OCT4/SOX2, and HNF1B, respectively, in cell lines of shared lineage where corresponding gene expression of TF>1 CPM. **L**. Pearson correlation coefficient (r) between TF gene expression (Z-score RPKM) and BAF occupancy as a function of corresponding TF motif archetype counts (Z-score(SMARCA4 ChIP-seq RPKM / [Motif])) across unique enhancer sites from B (C4-24) with r>0.5.

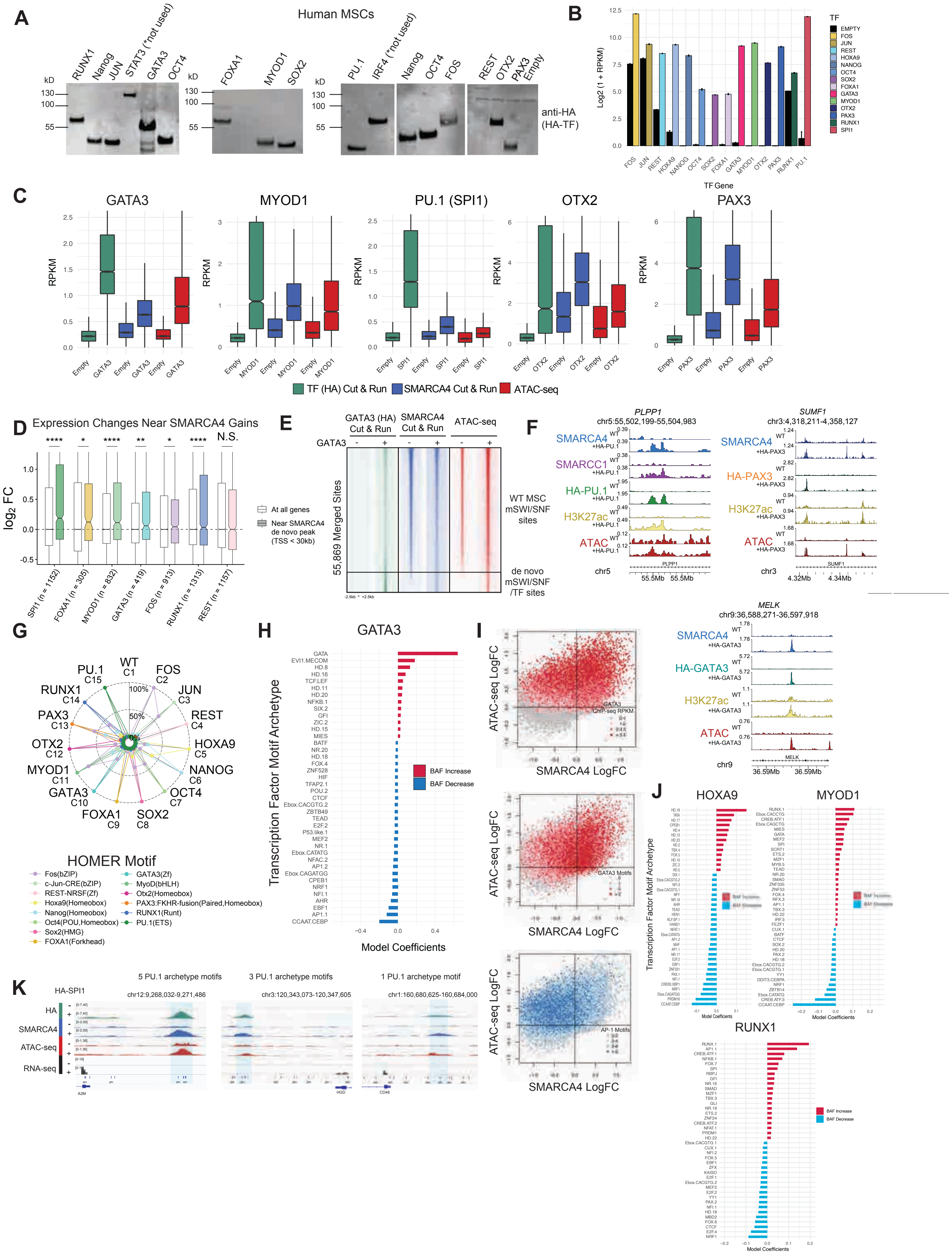

A
Human MSCs
RUNX1 Nanog JUN STAT3 (*not used) GATA3 OCT4
FOXA1 MYOD1 SOX2
PU.1 IRF4 (*not used) Nanog OCT4 FOS
REST OTX2 PAX3 Empty
kD
130
100
55
anti-HA (HA-TF)
B
Log2 (1 + RPKM)
TF Gene
TF
EMPTY
FOS
JUN
REST
HOXA9
NANOG
OCT4
SOX2
FOXA1
GATA3
MYOD1
OTX2
PAX3
RUNX1
SPI1
C
GATA3
MYOD1
PU.1 (SPI1)
OTX2
PAX3
RPKM
TF (HA) Cut & Run
SMARCA4 Cut & Run
ATAC-seq
D
Expression Changes Near SMARCA4 Gains
log2 FC
At all genes
Near SMARCA4 de novo peak (TSS < 30kb)
SPI1 (n = 1152)
FOXA1 (n = 305)
MYOD1 (n = 832)
GATA3 (n = 419)
FOS (n = 913)
RUNX1 (n = 1313)
REST (n = 1157)
N.S.
E
GATA3 (HA) Cut & Run
SMARCA4 Cut & Run
ATAC-seq
GATA3
55,869 Merged Sites
WT MSC mSWI/SNF sites
de novo mSWI/SNF /TF sites
F
PLPP1
chr5:55,502,199-55,504,983
SUMF1
chr3:4,318,211-4,358,127
SMARCA4
SMARCC1
HA-PU.1
H3K27ac
ATAC
HA-PAX3
WT
+HA-PU.1
+HA-PAX3
chr5
55.5Mb 55.5Mb
chr3
4.32Mb 4.34Mb
MELK
chr9:36,588,271-36,597,918
HA-GATA3
+HA-GATA3
chr9
36.59Mb 36.59Mb
G
WT C1
FOS C2
JUN C3
REST C4
HOXA9 C5
NANOG C6
OCT4 C7
SOX2 C8
FOXA1 C9
GATA3 C10
MYOD1 C11
OTX2 C12
PAX3 C13
RUNX1 C14
PU.1 C15
100%
50%
HOMER Motif
Fos(bZIP)
c-Jun-CRE(bZIP)
REST-NRSF(Zf)
Hoxa9(Homeobox)
Nanog(Homeobox)
Oct4(POU,Homeobox)
Sox2(HMG)
FOXA1(Forkhead)
GATA3(Zf)
MyoD(bHLH)
Otx2(Homeobox)
PAX3:FKHR-fusion(Paired,Homeobox)
RUNX1(Runt)
PU.1(ETS)
H
GATA3
Transcription Factor Motif Archetype
Model Coefficients
BAF Increase
BAF Decrease
I
ATAC-seq LogFC
SMARCA4 LogFC
GATA3 ChIP-seq RPKM
GATA3 Motifs
AP-1 Motifs
J
HOXA9
MYOD1
RUNX1
Transcription Factor Motif Archetype
Model Coefficients
BAF Increase
BAF Decrease
K
HA-SPI1
5 PU.1 archetype motifs
3 PU.1 archetype motifs
1 PU.1 archetype motif
chr12:9,268,032-9,271,486
chr3:120,343,073-120,347,605
chr1:160,680,625-160,684,000
HA
SMARCA4
ATAC-seq
RNA-seq
A2M
HGD
CD48

**Fig. S2: Ectopic expression of TFs direct mSWI/SNF genomic targeting and activity. B.** Immunoblots for HA-tagged TFs expressed in hMSCs (whole cell extracts). **B.** Expression (Log(1+RPKM)) of TFs overexpressed in MSCs relative to their endogenous level of expression in cells transduced with empty vector. **C.** Enrichment (RPKM) of HA-tagged TF, SMARCA4, and DNA accessibility (ATAC-Seq) at TF-specific peaks (as labeled above each plot) in hMSCs expressing empty vector or TFs indicated on X-axis. **D.** Log2FC gene expression changes upon TF expression (relative to empty control) of all genes (shown in white) versus genes near de novo SMARCA4 peaks (shown in colors). Significance indicated above each TF comparison. **E.** Occupancy (RPKM normalized) profiles of HA-tagged GATA3, SMARCA4 and DNA accessibility at 55,869 mSWI/SNF sites in MSCs expressing GATA3 or empty vector control. Wild-type mSWI/SNF sites are indicated on top and de novo GATA3-dependent peaks on the bottom. **F.** Example CUT&RUN tracks of HA-tagged PU.1, GATA3, PAX3, as well as SMARCA4, H3K27Ac, DNA accessibility (ATAC-Seq) and gene expression (RNA-Seq) at selected loci. **G.** Homer motif enrichment ($\log_{10}$p-value) of indicated TFs in peak clusters defined in Figure 1B. **H.** GLMnet motif enrichment analysis identifies top TF motifs underlying SMARCA4 peaks that displayed gain of SMARCA4 enrichment (in red) and loss of enrichment (in blue) upon TF overexpression. **I.** Scatterplots displaying the correlation between change in SMARCA4 occupancy (x-axis) and DNA accessibility measured by ATAC-Seq (Y-axis) in MSCs expressing PU.1 compared to empty vector. Color key indicates PU.1 RPKM enrichment (top panel), number of PU.1 motifs (middle panel) and number of AP.1 motifs (lower panel). **J.** GLMnet motif enrichment identifies top TF motifs underlying SMARCA4 peaks that displayed gain of SMARCA4 enrichment (in red) and loss of enrichment (in blue) upon TF overexpression. **K.** Example CUT&RUN tracks for HA-tagged PU.1, SMARCA4, DNA accessibility (ATAC-Seq) and gene expression (RNA-Seq) at loci containing 5, 3 and 1 PU.1 archetypical motifs, respectively.

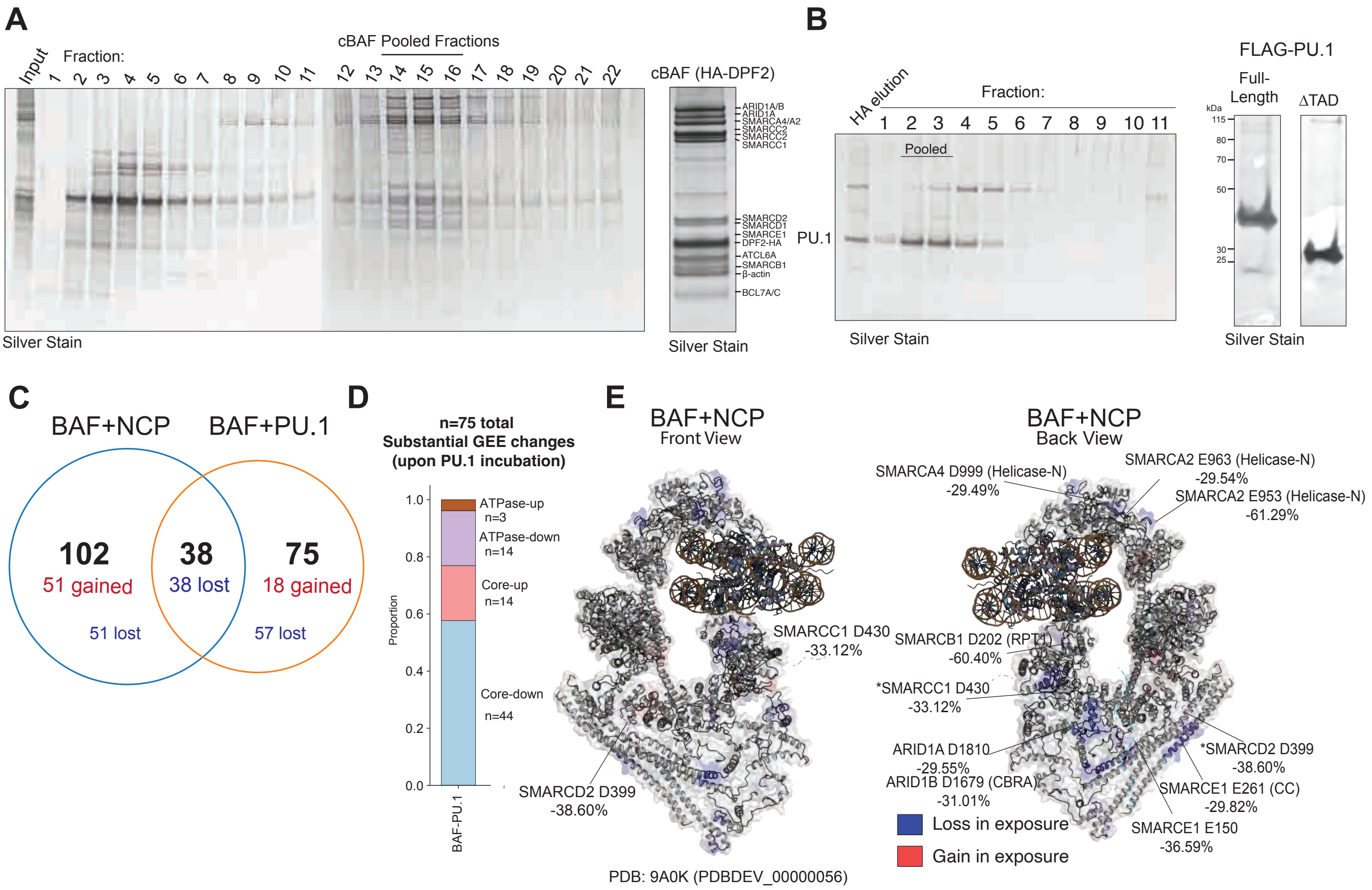


**Fig. S3: Protein footprinting experiments identify a putative PU.1 interaction interface on the human cBAF complex. A.** Purification of endogeneous cBAF complexes from HEK-293T cells using HA affinity purification of HA-tagged DPF2 subunit, followed by density sedimentation. Silver stains for cBAF gradient fractions (left) and pooled fractions (Fx 14-15, right)**. B.** Silver stained SDS-PAGE gel showing purification of PU.1 using HA affinity purification followed by density gradient sedimentation used in GEE labeling experiments. **C.** Venn diagram showing overlap between peptides exhibiting changes in GEE labeling of solvent-exposed residues upon incubation with NCP or PU.1. **D.** Distribution of cBAF peptides differentially labeled upon PU.1 interaction within ATPase or Core module subunits. **E.** cBAF peptides with detectable changes in GEE labeling upon NCP binding are mapped on the cBAF-NCP 3D structure (PDB: 9A0K(PDBDEV_00000056)).

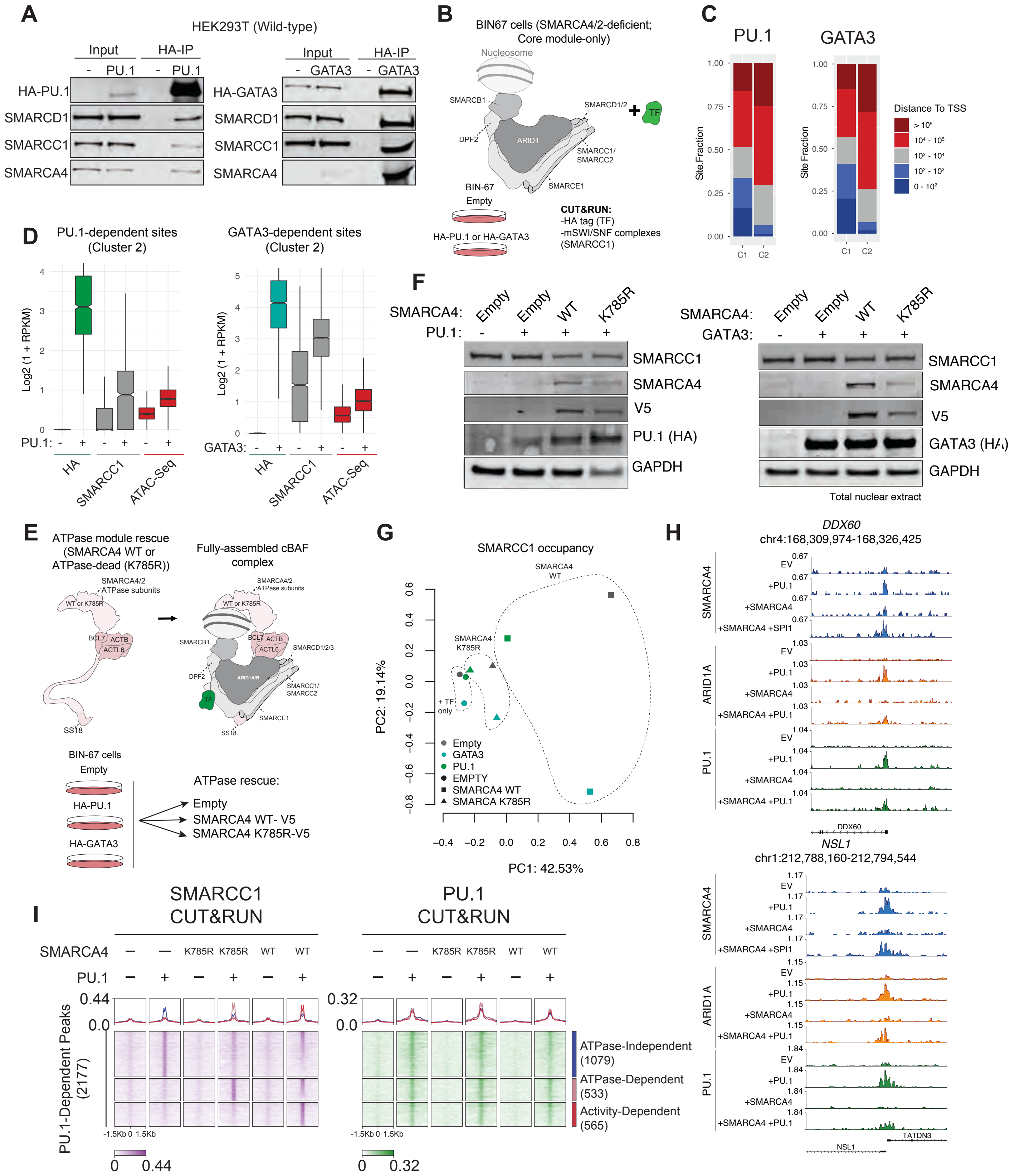


**Fig. S4: The cBAF core is sufficient for PU.1 and GATA3 TF interaction and mSWI/SNF chromatin targeting in cells. A.** Immunoblots performed on immunoprecipitation studies of HA-tagged PU.1 and GATA3 expressed in wild-type HEK293T cells. **B.** Schematic for experiments in which HA-tagged PU.1 or GATA3 were expressed in SMARCA2/SMARCA4-deficient BIN67 SCCOHT cells and occupancy of mSWI/SNF core module subunits were probed using CUT&RUN. **C.** Distance-to-TSS stacked bar graphs quantifying the occupancies of HA-tagged TF, SMARCC1 and DNA accessibility at TF-dependent sites in BIN67 cells expressing PU.1 or GATA3. **D.** Occupancy of HA-tagged TF and SMARCC1, ATAC-seq over TF-dependent sites (Cluster 2). **E.** Schematic for experiments rescuing SMARCA4 WT or ATPase dead (K785R) relative to empty vector control in BIN67 SSCCOHT cells with and without concomitant expression of TFs. **F.** Expression of PU.1 or GATA3 TFs with or without concomitant rescue of SMARCA4 WT or ATPase-dead variants. **G.** PCA analysis of SMARCC1 occupancy across conditions indicated. **H.** Representative tracks at the *DDX80* and *NSL1* loci depicting TF-mediated redirection of the mSWI/SNF core module in BIN67 cells. **I.** Heatmaps for SMARCC1 and PU.1 (SPI1) over PU.1-dependent peaks. ATPase-independent, ATPase-dependent, and full complex (activity)- dependent sets of sites are indicated.

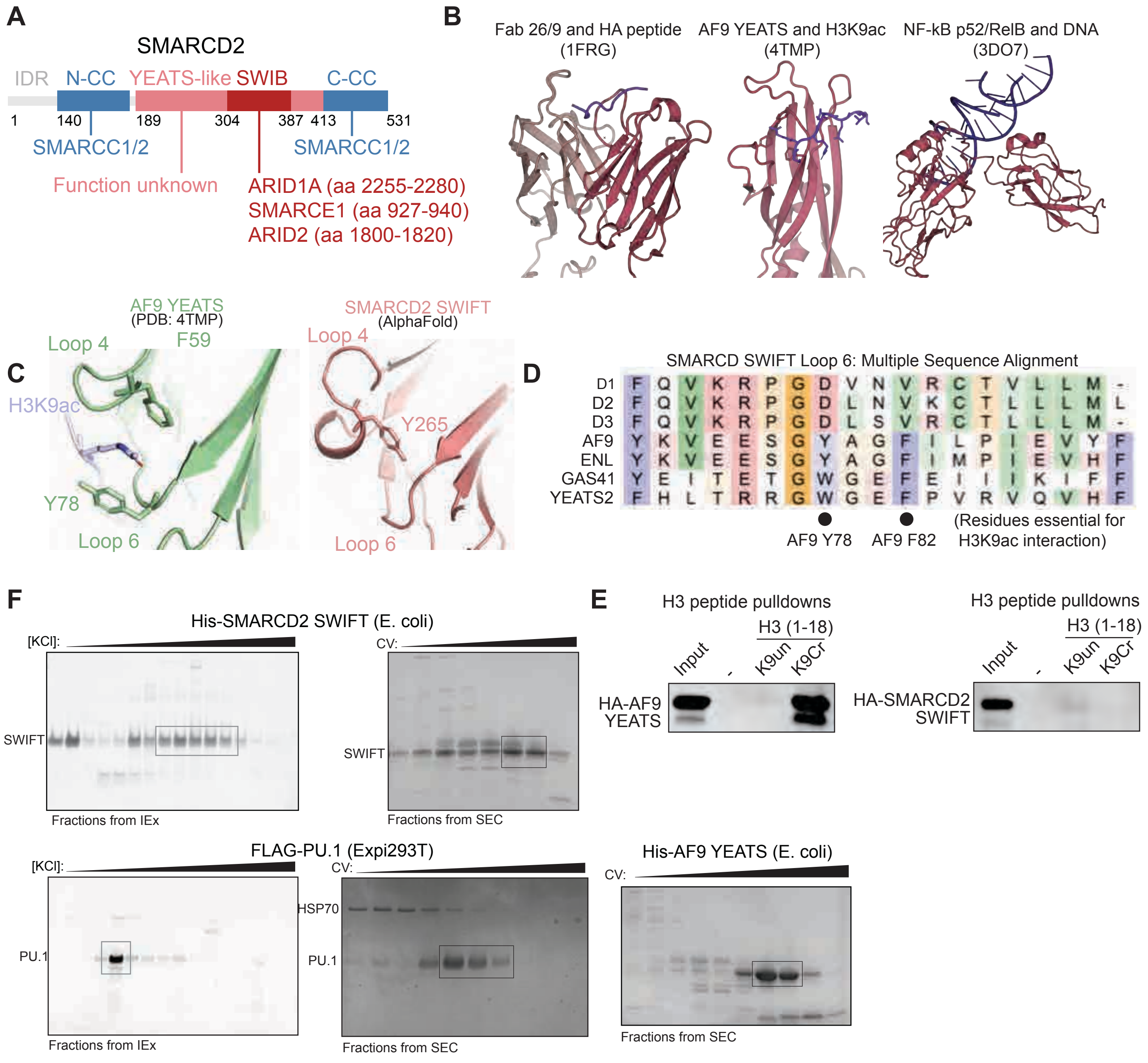


**Fig. S5: Characterization of the SMARCD Ig-fold-like SWIFT domain. A.** Domain structure of the human SMARCD2 subunit with known binding interactions within mSWI/SNF complexes indicated. The coiled-coil domains of SMARCD subunits (blue) make extensive interactions with SMARCC1/2 subunits; SWIB interacts with cBAF-specific ARID1A or PBAF-specific ARID2 subunits. YEATS-like Ig domain does not engage with other mSWI/SNF subunits and its function is unknown. **B.** Structures of immunoglobulin domains and their diverse interaction partners; (*from left to right*) IgG with HA peptide, AF9 YEATS domain with H3K9ac peptide, NFkB bound to its cognate DNA. **C.** Structural comparison of AF9 H3K9ac-binding aromatic tunnel formed between loop 4 and loop 6. Essential F59 and Y78 residues engage in $\pi$-$\pi$-$\pi$ interactions with the H3K9ac group. **D.** Multiple sequence alignment of loop 6 of human SMARCD1/2/3 SWIFT domains with YEATS domains from AF9, ENL, GAS41 and YEATS2. The essential resides (Y78 and F82) for H3K9ac interaction are absent from the SWIFT domain loop 6, suggesting a functional divergence. **E.** Peptide pulldowns of AF9 YEATS domain (left) and SMARCD2 SWIFT domain (right) with histone H3 (aa1-18) K9unmodified or H3 (aa1-18) K9crotonylated peptides. **F.** Purification of SWIFT domain from E. coli and PU.1 from Expi293F cells using anion exchange followed by size exclusion chromatography for in vitro binding studies.

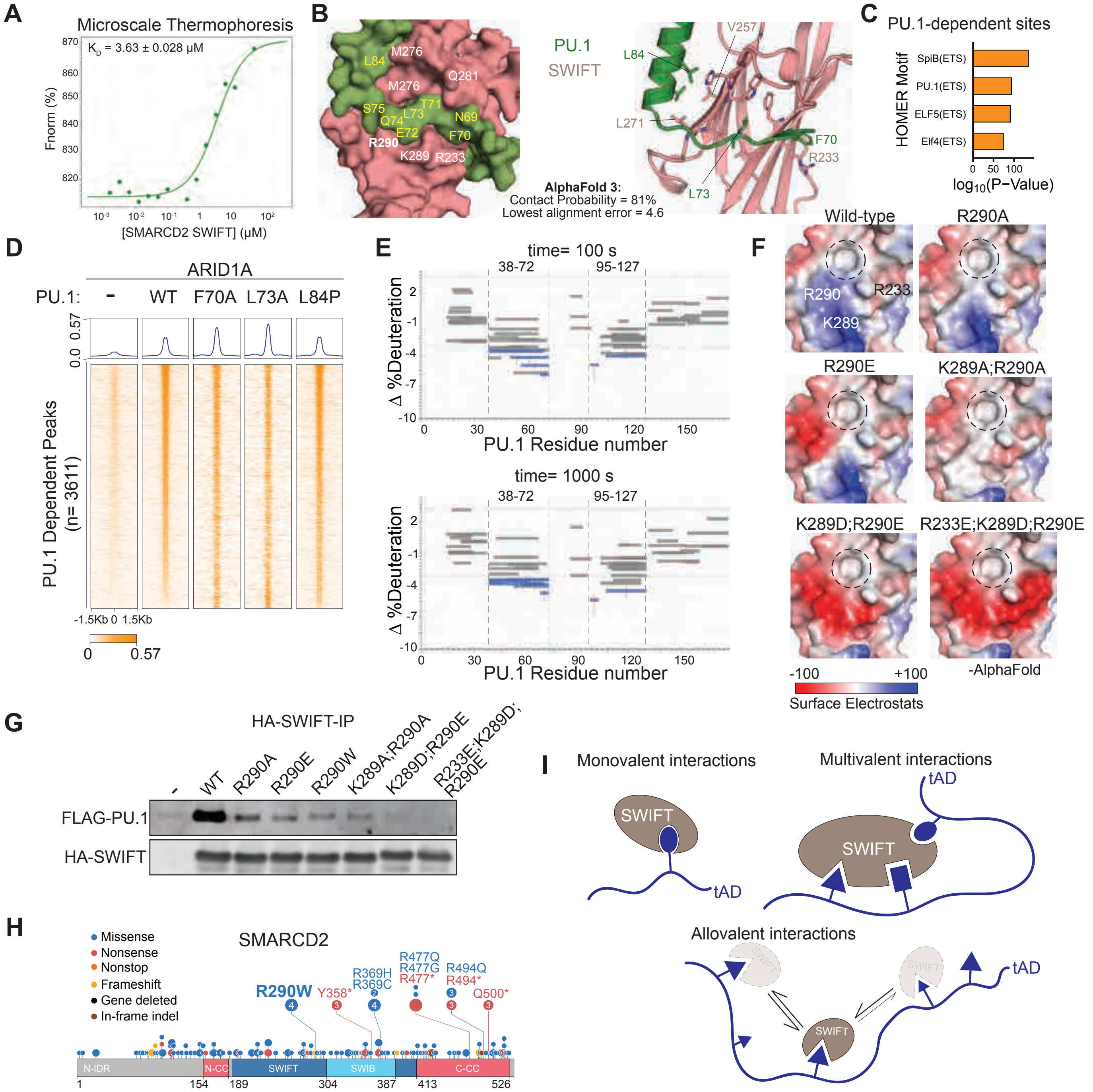


**Fig. S6: Characterization of the interaction between PU.1 and the SMARCD2 SWIFT domain. A.** $K_d$ model fit for interaction between SMARCD2 SWIFT domain and PU.1 using microscale thermophoresis (Kd = 3.6uM). **B.** AlphaFold multimer modeling of SMARCD2 SWIFT domain with PU.1 showing a interaction between PU.1 L73 and hydrophobic pocket residues on the SWIFT domain. Space filling model of interaction interface is shown on the right, displaying the proximity of electronegative and acidic residues to K289 and R290 basic residues. **C.** HOMER motif analysis over PU.1-dependent sites in hMSCs. **D.** Heatmaps showing CUT&RUN occupancy of ARID1A in hMSCs expressing PU.1 WT and mutant variants. **E.** Change in % Deuterium uptake by each PU.1 peptide upon addition of SMARCD2 SWIFT domain. Dashed line represent the threshold of statistically significant changes. Error bars represent standard deviation. Regions of signal are indicated; t= 100s (top), t=1000s (bottom). **F.** Surface potential of the SMARCD2 SWIFT domain with indicated point mutation(s). **G.** Immunoblots of HA-SWIFT immunoprecipitations from Expi293F cells expressing FLAG-tagged PU.1 and indicated HA-tagged SWIFT mutants. **H.** Cancer-associated mutations found in COSMIC database are shown on the domain structure of SMARCD2. **I.** Schematic depicting non-mutually exclusive modes of protein-protein interactions. In monovalent interactions, single binding sites on tADs interact with a single receptor site on SWIFT. In multivalent interactions, multiple specific binding sites on tAD of TFs interact with distinct residues on SWIFT in a particular configuration. In an allovalent mode of interaction, multiple binding motifs on a tAD interact with the same pocket on SWIFT with varying affinities. An equilibrium between various SWIFT binding states increases the overall dwelling time of SWIFT on the tAD, increasing the observed overall association constant between SWIFT and TF. Allovalent interactions are expected to require a shared “grammar” of amino acid composition on tADs, but not a specific sequence conservation within tADs of various TFs.

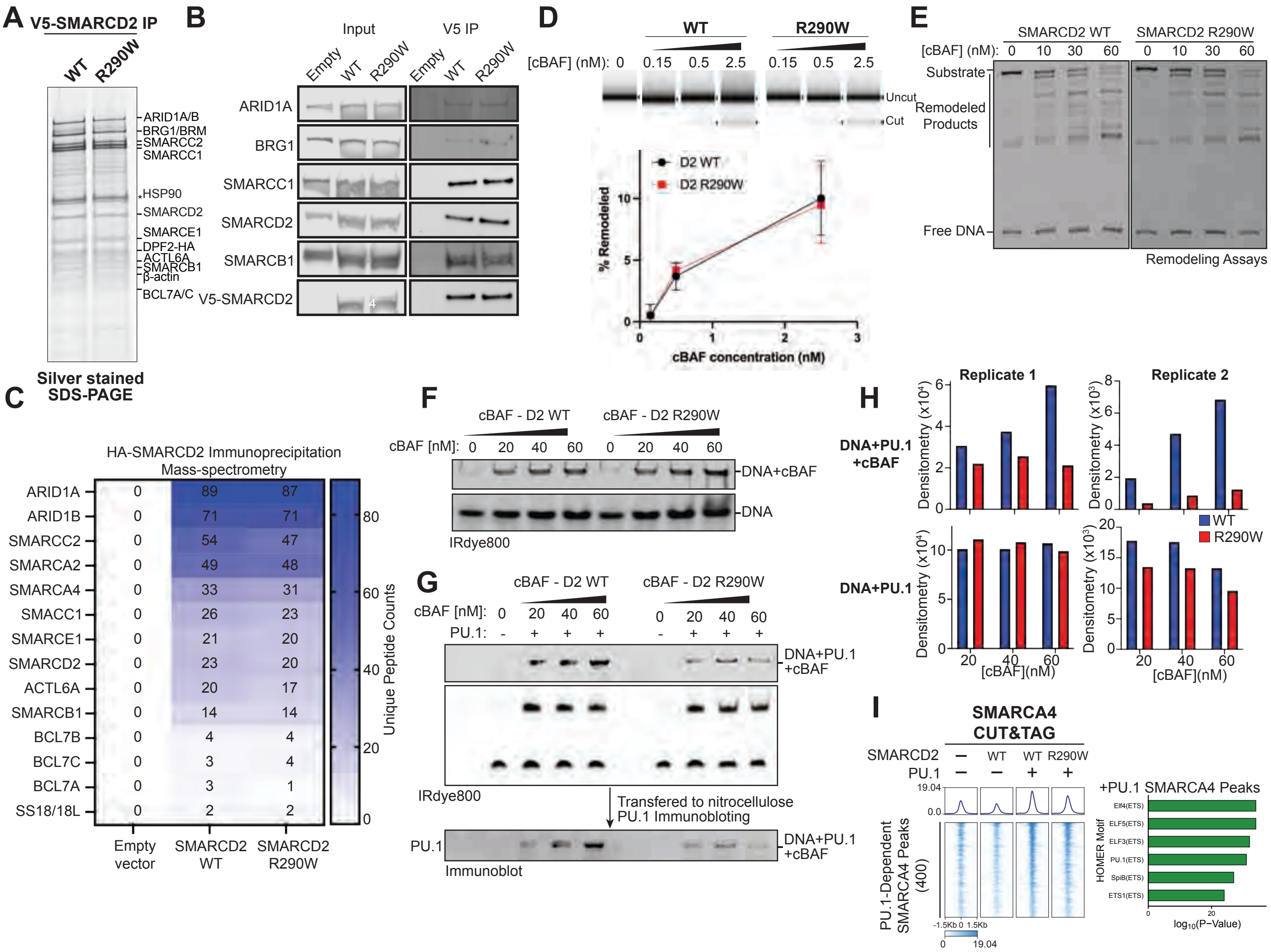


**Fig. S7: SMARCD2 containing the R290W SWIFT point mutation assembles into enzymatically active mSWI/SNF complexes. A.** Silver stained SDS-PAGE showing the subunits of mSWI/SNF complexes co-purified with V5-tagged SMARCD2 WT and R290W mutant from HEK-293T cells. **B.** Immunoblots of select mSWI/SNF complex subunits immunoprecipitated with V5-tagged SMARCD2 WT or R290W mutant complexes. **C.** Proteomic mass spectrometric analysis of mSWI/SNF complex subunits purified with V5-tagged SMARCD2 WT or R290W mutant. **D.** REAA assays performed with 5 nM nucleosomes incubated with varying concentrations of mSWI/SNF complexes containing SMARCD2 WT or R290W mutant, ATP and DpnII restriction enzyme for 30 min. Quantification of remodeled products as measured DNA size distribution is shown below. **E.** Nucleosome sliding assays using 50 nM nucleosomes incubated with varying concentrations of mSWI/SNF containing SMARCD2 WT or R290W mutant for 30 min. Products were visualized using native PAGE gel. **F.** Electromobility Shift Assays (EMSA) showing intrinsic DNA binding affinities of cBAF complexes containing SMARCD2 WT or R290W. **G.** EMSA to assess binding between cBAF and DNA probe in the presence of PU.1 factor. Following electrophoresis, products were transferred to a nitrocellulose membrane and blotted with PU.1 antibody. **H.** Densitometry quantification of cBAF-PU.1-DNA (top) and PU.1-bound DNA (bottom) ternary complexes from EMSA. Intensities of the free DNA band was outside the linear range of quantification. **I.** SMARCA4 CUT&TAG performed in hMSCs infected with empty vector, WT SMARCD2, or R290W SMARCD2, with or without PU.1. *Left*, heatmap over PU.1-dependent peaks; *Right*, HOMER motif analysis over peaks.

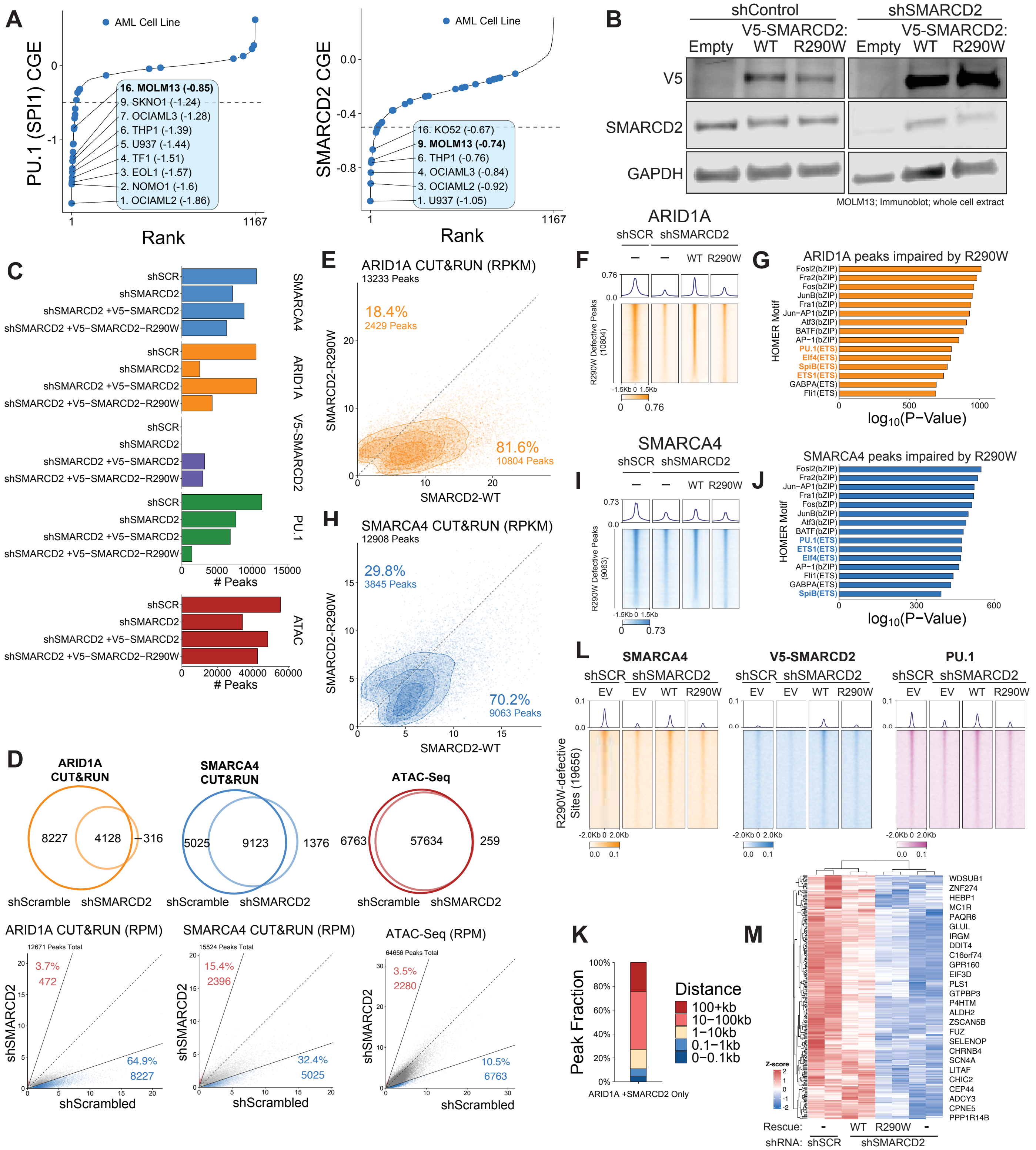


**Fig. S8: SWIFT-mediated interaction between cBAF and PU.1 is necessary for genomic targeting and activity of mSWI/SNF in MOLM-13 AML cells. A.** 1167 cell lines are ranked by their CRISPR dependency score (CRISPR gene effect, CGE) of PU.1 (left) or SMARCD2 (right). AML cell lines are marked in blue circles. **B.** Immunoblots of whole cell extracts from MOLM-13 cells expressing shRNA to target endogenous SMARCD2 or a non-targeting control, and rescued with an shRNA-resistant SMARCD2 WT or R290W transgene. **C.** Number of SMARCA4, ARID1A, V5-tagged SMARCD2 and PU.1 CUT&RUN peaks identified in SMARCD2 knockdown MOLM13 cells rescued with SMARCD2 WT or R290W transgenes. Number of ATAC-Seq peaks are shown below in red. **D.** Overlap between ARID1A, SMARCA4 and ATAC-Seq peaks in MOLM13 cell expressing shRNA against SMARCD2 or a non-targeting control. **E.** Scatterplot showing ARID1A CUT&RUN RPM enrichment in SMARCD2-depleted MOLM13 cells rescued with SMARCD2 WT or R290W mutant. **F.** Heatmaps displaying the ARID1A occupancies in MOLM13 cells at 81.6% of genomic sites that showed impaired targeting. **G.** HOMER motif enrichment analyses of ARID1A peaks with reduced occupancy in cells expressing SMARCD2 R290W. **H-J.** Same as E-G but for SMARCA4 CUT&RUN. **K.** Stacked bar chart showing the distance from transcription start sites of the ARID1A peaks recovered by re-expression of SMARCD2 WT, but not R290W mutant. **L.** Heatmaps showing SMARCA4, V5-tagged SMARCD2 and PU.1 occupancy profiles at cBAF-occupied sites in MOLM-13 cells rescued with empty vector (EV), WT SMARCD2, or R290W mutant following shRNA-mediated endogenous SMARCD2 suppression. **M.** Heatmap showing Z-score normalized gene expression RPKM of selected known PU.1-target genes in conditions indicated.

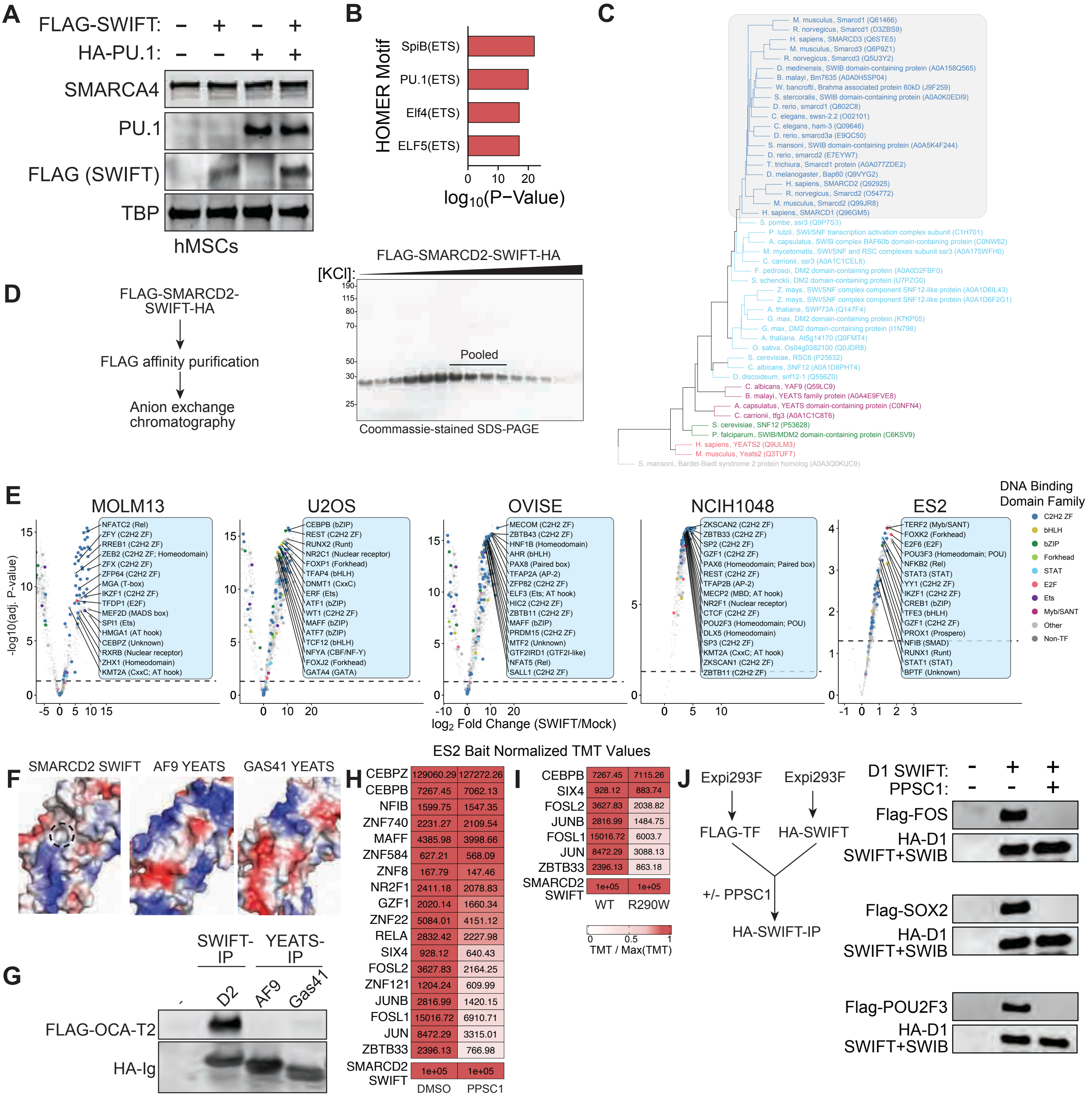


**Fig. S9: SWIFT is a transcription factor interaction platform for mSWI/SNF. A.** Immunoblots of whole cell extracts from hMSCs overexpressing PU.1 in the presence or absence of ectopic expression of FLAG-tagged SMARCD2 SWIFT domain. **B.** HOMER motif enrichment analysis of PU.1-dependent de novo SMARCA4 genomic target loci. **C.** Phylogenetic analysis of SWIFT domain primary sequence reveals its evolutionary conservation throughout the phylogenetic tree. **D.** Purification of N-terminally FLAG-tagged and C-terminally HA-tagged SMARCD2 SWIFT domain from mammalian Expi293F cells using FLAG affinity purification. **E.** Scatterplots displaying the enrichment of TFs bound by 200 nM SWIFT domain incubated with ammonium sulfate-extracted nuclear lysates from n=5 human cancer lines spanning diverse tissue types. **F.** Comparison of surface potential of SMARCD2 SWIFT and the YEATS domains of AF9 and GAS41. **G.** HA immunoprecipitation from cells co-expressing FLAG-tagged OCA-T2 with HA-tagged SMARCD2 SWIFT domain or YEATS domains of AF9 and GAS41. **H.** TMT peptide enrichment in ES2 cells (containing isolated SWIFT) but in the presence or absence of 50 μM PPSC1. **I.** Same as H but with SWIFT WT or R290W. **J.** HA-SWIFT immunoprecipitation from cells co-expressing SMARCD1 SWIFT domain with the indicated TF in the presence of absence of 50 μM PPSC1.

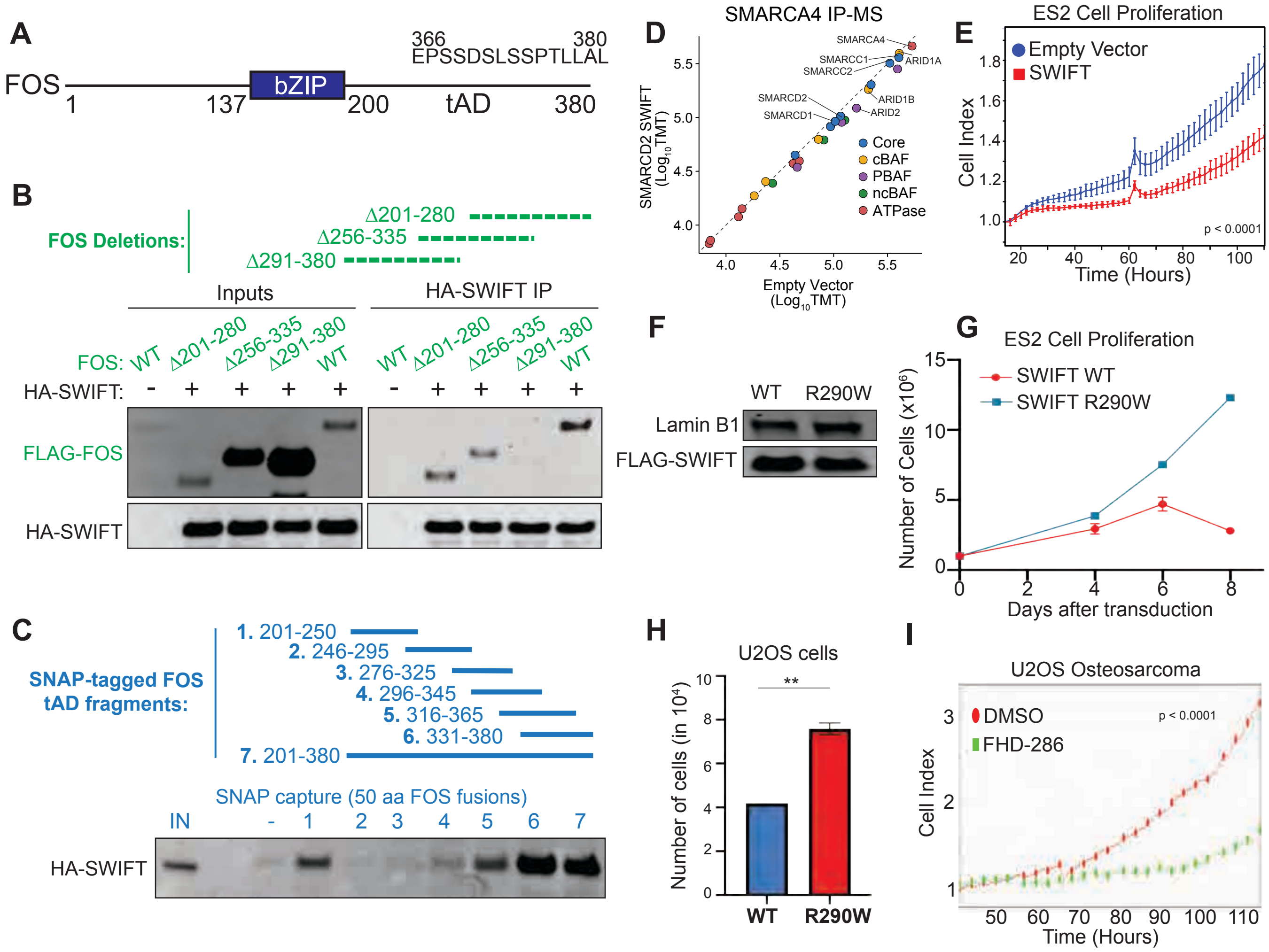


**Fig. S10: Dominant expression of the SWIFT domain impairs cancer cell proliferation by sequestering endogenous mSWI/SNF from TFs. A.** Domain structure of the FOS transcription factor showing the bZIP DNA-binding domain and the C-terminal tAD. Leucine/Serine-rich motif is shown on top. **B.** Immunoblots of HA-SWIFT immunoprecipitation from cells co-expressing HA-tagged SMARCD1 SWIFT and indicated FLAG-tagged FOS deletion mutants. Dashed lines show the regions of the FOS tAD deletions. **C.** Immunoblots of eluates from pulldowns of SNAP-tagged FOS fragments with purified HA-tagged SMARCD1 SWIFT domain. **D.** Scatterplot showing the enrichment of mSWI/SNF complex subunits bound to SMARCA4 from U2OS cells expressing empty vector or HA-tagged SMARCD2 SWIFT domain, confirming complex integrity. **E.** Proliferation of U2OS cells overexpressing SWIFT domains, monitored by eSight. P-value $p<0.0001$ of difference in the slope of exponential growth for each cell line (n= 3 replicates) is indicated. **F.** Immunoblots of ES2 cells expressing FLAG-tagged WT or R290W mutant SMARCD2 SWIFT domain. **G.** Proliferation of ES2 cells overexpressing FLAG-tagged WT or R290W mutated SWIFT domain. **H.** Number of U2OS cells on day 6 after expression of the SWIFT domain of SMARCD2 WT or R290W. p-value of 0.046 was determined by t-test **I.** Proliferation of U2OS cells treated with 100 nM FHD-286 or DMSO control (n= 2 replicates). P-value $p<0.0001$ of difference in the slope of exponential growth for each cell line is indicated.

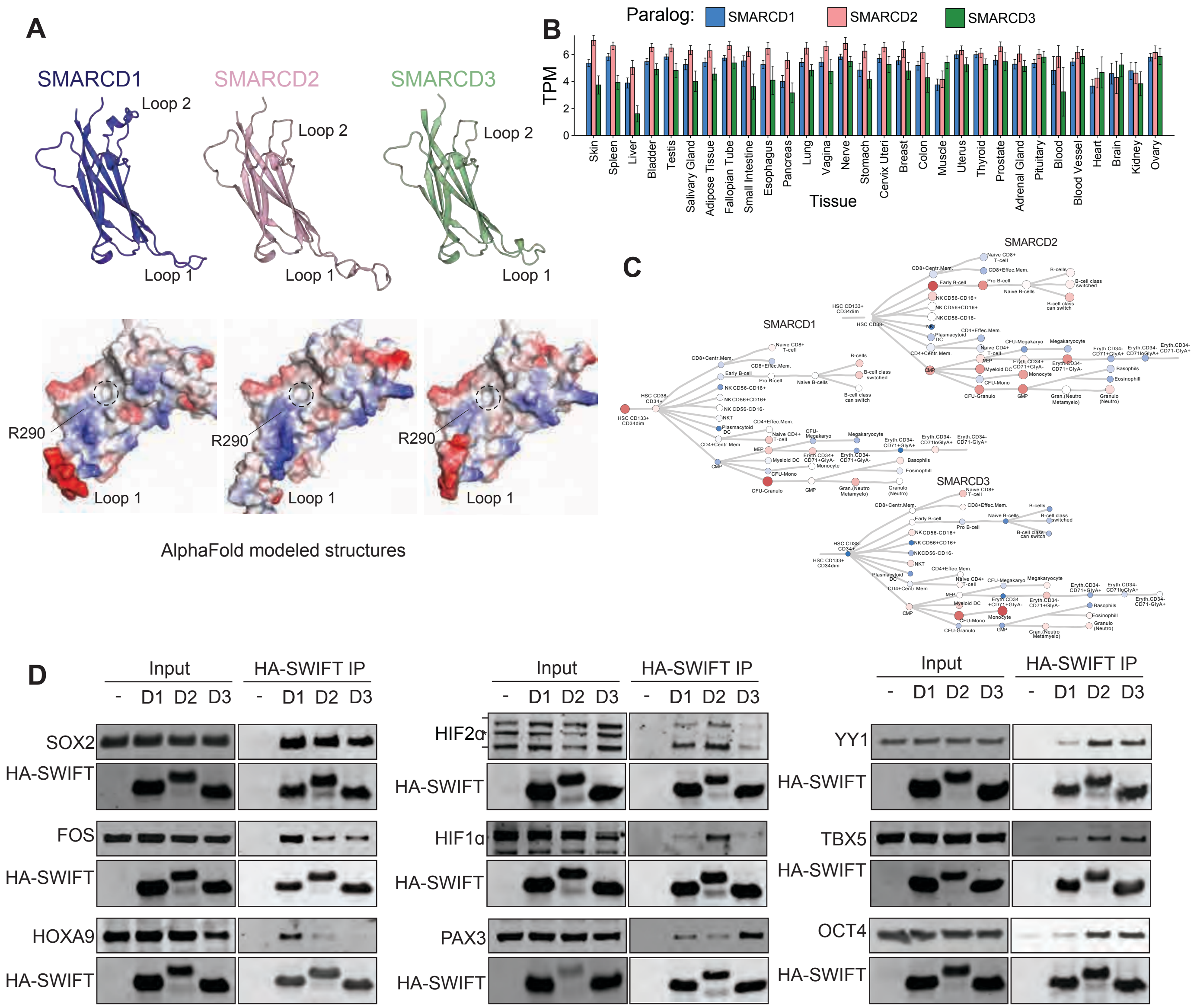


**Fig. S11: Transcription factors preferentially interact with SWIFT domains of specific SMARCD paralogs. A.** Structural predictions of SMARCD1, SMARCD2 and SMARCD3 SWIFT domains using AlphaFold. **B.** Barcharts showing the expression of SMARCD paralogs in various human tissues. **C.** Expression of SMARCD1, SMARCD2 or SMARCD3 paralogs in various cell types of the hematopoietic lineage. **D.** Immunoblots of HA-SWIFT immunoprecipitation from HEK293T cells co-expressing HA-tagged SWIFT of specific SMARCD paralogs (D1, D2, D3) and the indicated FLAG-tagged transcription factors.

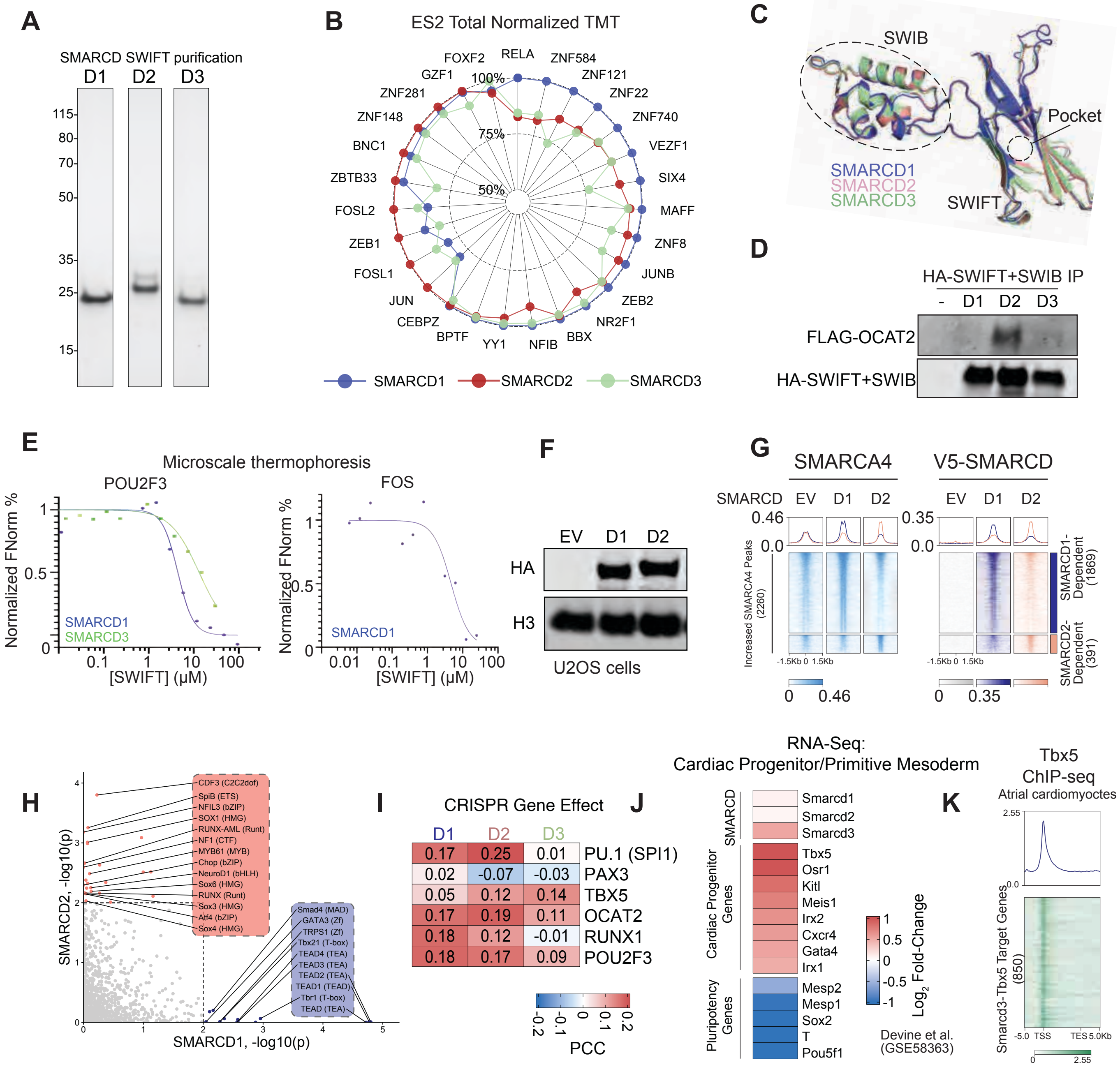


**Fig. S12: Transcription factors preferentially interact with SWIFT domains of specific SMARCD paralog. A.** Coomassie-stained SDS-PAGE gels of N-terminally His-tagged and C-terminally HA-tagged SWIFT domains of SMARCD1, SMARCD2 and SMARCD3 paralogs purified from E.coli. **B.** Radar plot showing the preferences of TFs for the SWIFT domains of SMARCD1 (blue), SMARCD2 (red), SMARCD3 (green) paralogs. **C.** Alphafold predicted 3D structure of SMARCD1, D2, and D3 SWIFT domains along with the SWIB domain within its loop 7. **D.** Immunoblots of HA-SWIFT immunoprecipitation from HEK293T cells expressing FLAG-tagged OCA-T2 and HA-tagged SWIFT+SWIB of SMARCD1/2/3 paralogs. **E.** Microscale thermophoresis to identify dissociation constants of interactions between POU2F3 and FOS and SMARCD1/2/3 SWIFT domains. Hill curve was fitted on normalized Fnorm values for data visualization. Dissociation constants shown in Fig. 6E were determined by fitting a Kd model for one-site binding. **F.** Immunoblots of U2OS cells expressing HA-tagged SMARCD1 or SMARCD2. **G.** Heatmaps displaying the enrichment of HA-tagged SMARCD1 or SMARCD2 at their de novo peaks. **H.** Scatter plot showing motif enrichment for SMARCD1- and SMARCD2-specific peaks, as defined in **G. I.** Heatmap showing correlation between CRISPR gene effect of SMARCD paralogs (D1, D2, D3) and lineage-specific TFs in Fig. 6H. **J.** Heatmap showing the gene expression pattern (z-score normalized FPKM) of co-expressed TFs during cardiomyocyte differentiation. **K.** Heatmap of TBX5 ChIP-Seq profile at gene bodies of SMARCD3-TBX5 co-regulated genes.

**Table S1. Antibodies used in this study**

| Antibody | Vendor | Catalog# | Lot Number | Application |
|---|---|---|---|---|
| HA epitope | Cell Signaling | 3724 | monoclonal | CUT&RUN, Immunobloting |
| FLAG epitope | Cell Signaling | 2368 | monoclonal | immunobloting |
| SMARCA4 | Cell Signaling | 49360 | monoclonal | CUT&RUN, Immunobloting |
| PU.1 | Cell Signaling | 2258 | monoclonal | CUT&RUN, Immunobloting |
| ARID1A | Cell Signaling | 12354 | monoclonal | CUT&RUN, Immunobloting |
| SMARCC1 | Cell Signaling | 11956 | monoclonal | CUT&RUN, Immunobloting |
| SMARCD1 | Cell Signaling | 35070 | monoclonal | Immunobloting |
| IgG | Epicypher | 13-0042 | monoclonal | CUT&RUN, Immunobloting, Immunoprecipitation |
| H3K27ac | Cell Signaling | 8173 | monoclonal | CUT&RUN |
| V5 | Cell Signaling | 13202 | monoclonal | CUT&RUN, Immunobloting |
| SMARCA4 | Abcam | ab110641 | monoclonal | Immunoprecipitation |

**Table S2. Cell lines used in this study**

| Cell Line | Species | RRID | Vendor | Catalog # |
|---|---|---|---|---|
| MOLM13 | human | CVCL_2119 | | |
| Lenti X 293T | human | | Takara | 632180 |
| Expi293F | human | | Thermo | A14527 |
| U2OS | human | CVCL_0042 | | |
| ES2 | human | CVCL_AX39 | | |
| Ovise | human | CVCL_3116 | | |
| hMSASC52telo | human | CVCL_U602 | ATCC | SCRC-4000 |
| NCI-H1048 | human | CVCL_1453 | | |

**Table S3. Sequences of oligos for generating EMSA probes**

| | Forward | Reverse |
|---|---|---|
| Control (TF non-binding) | ATGCAATAAGATACGTTCTAGAGAGGTTACGATAG | CTATCGTAACCTCTCTAGAACGTATCTTATTGCAT |
| PU.1 binding probe | TGAAATAACCTCTGAAAGAGGAACTTGGTTAGGTA | TACCTAACCAAGTTCCTCTTTCAGAGGTTATTTCA |

**Supplementary Text**

Modes of protein-protein interactions

Ligand-receptor interactions can vary in complexity depending on the number and arrangement of binding sites, giving rise to mono-, multi-, and allovalent binding modes (**Fig. S6I**). These distinct binding modes influence the strength and dynamics of molecular interaction. Therefore, we define these key concepts below:

**Monovalency** refers to an interaction in which a molecule, such as a ligand or protein domain, engages a single binding site on its target. This one-to-one interaction typically exhibits defined stoichiometry and affinity.

**Multivalency** describes the capacity of a molecule or complex to engage multiple binding sites simultaneously, either on the same target (intra-molecular) or on different targets (inter-molecular). Multivalent interactions can increase overall binding strength (avidity) and specificity through cooperative or cumulative effects of individual low-affinity contacts.

**Allovalency** is a special case of multivalency in which a single ligand possesses multiple distinct binding sites that can independently engage a target through alternative, mutually exclusive interactions. In allovalent systems, only one site binds at a time, but the presence of several potential binding motifs increases the effective affinity by enhancing the probability of productive engagement. A common example of allovalency is: affinity of 3X-FLAG peptide (with slight variation in the individual linear sequence) for FLAG antibody exceeds that of a 1X FLAG sequence. This is despite the fact that at a time, only one FLAG sequence within the 3X-FLAG can engage with the antibody. However, the presence of three 'allovalent' sites increases the overall apparent affinity between the two ligands.

Electromobility Shift Assays

**Complex Quality Controls:** To ensure equal complex loading, we quantified purified cBAF complex concentrations using both Nanodrop and Qubit measurements and validated these measurements by western blot quantification of BRG1 using purified BRG1 protein as a standard (Epicypher, SKU: 15-1014). We further confirmed equal total protein amounts across preparations by SDS-PAGE followed by silver staining and immunoblotting (**Fig. S7A-B**). To verify that all complexes shared equivalent subunit composition, we performed mass spectrometry analysis, which ruled out stoichiometric variation as a potential source of the observed differences in electrophoretic mobility (**Fig. S7B-C**). We next assessed the biochemical activity of each purified complex by performing nucleosome sliding and restriction enzyme (DpnII) accessibility assays. These experiments confirmed that the SMARCD2 R290W mutation does not alter the intrinsic remodeling activity of cBAF complexes (**Fig. S7D-E**). The equal biochemical activities observed in vitro further support equal complex loading across preparations.

**EMSA Controls:** To rule out differences in DNA-binding ability as a source of the observed differences in cBAF-dependent DNA mobility shifts, we performed electrophoretic mobility shift assays (EMSAs) with increasing concentrations of cBAF alone, in the absence of PU.1 (**Fig. S7F**). Both SMARCD2 WT- and R290W- containing complexes exhibited equivalent DNA binding under these conditions, indicating that the altered supershift upon PU.1 addition reflects changes in PU.1-cBAF interaction rather than nonspecific DNA binding (**Fig. S7F**). Notably, all EMSA experiments shown in **Fig. 3J** and **Fig. S7F-G** were performed in the presence of a 5X excess of unlabeled scrambled control probe to suppress nonspecific binding by SMARCA4 or other DNA-binding subunits within mSWI/SNF complexes.

Due to the large size of cBAF complexes (1.2 MDa) relative to PU.1 (35 KDa) and PU.1 antibody (~150 KDa), detection of PU.1 in cBAF-PU.1-DNA ternary complex is difficult by supershift assays. Therefore, to confirm that the cBAF-dependent super-shifted band contained PU.1, we transferred EMSA gels to nitrocellulose membranes and immunoblotted with PU.1 antibody. PU.1 was indeed detected within the cBAF-dependent super-shifted bands, consistent with the presence of a DNA-PU.1-cBAF ternary complex (**Fig. S7G**). Quantification of free DNA, DNA-PU.1, and DNA-PU.1–cBAF bands is provided in **Fig. S7I**. Because full-length cBAF complexes could not be concentrated to the micromolar range required for precise $K_D$ determination, we refrained from drawing conclusions regarding the affinity of assembled cBAF complexes for PU.1 from these experiment.

<u>Supplementary Methods:</u>

**SNAP capture affinity precipitation:** Expi293F cells (30 mL cultures) were transfected with 20 μg of plasmid DNA containing SNAP-tagged TF fragments using 5X PEI reagent. The following day, valproic acid, sodium propionate, 1X GlutaMax, and 1X nonessential amino acids (NEAA) were added to enhance protein expression. Cells were harvested on day 5 by centrifugation at 1,000 × g for 5 min, washed once with PBS, and resuspended in 4 mL of lysis buffer (20 mM HEPES pH 8.0, 200 mM KCl, 1 mM EDTA, 1× protease inhibitor cocktail, 1 mM DTT, 0.8 mM PMSF, and 0.01% NP-40). Lysates were mixed thoroughly by pipetting and incubated for ~1 h at 4°C. Two milliliters of extract were clarified by centrifugation at maximum speed for 15 min to remove chromatin.

For affinity capture, 80 μL of SNAP-capture beads (per reaction) were washed twice with lysis buffer and resuspended at 50 μL per reaction. Clarified lysates were transferred to fresh tubes and incubated with 50 μL of pre-equilibrated SNAP-capture beads for 3 h at room temperature with rotation. Beads were then washed 5 times with lysis buffer followed by two washes with binding buffer (20 mM HEPES pH 8.0, 75 mM KCl, 0.5 mM EDTA, 1 mM DTT, 1× protease inhibitor cocktail, and 0.01% Tween-20).

Beads from each reaction were resuspended in 20 μL of binding buffer and incubated with recombinant SMARCD1 SWIFT domain at a final concentration of 10 μM in binding buffer, overnight at 4°C with rotation; an aliquot of the master mix was saved as input control.

The following day, beads were separated on a magnetic rack, and supernatants were discarded. Beads were washed five times with binding buffer, using gentle pipette mixing at room temperature for 2 min per wash. After three washes, beads were transferred to new tubes, and the final two washes were performed in fresh tubes to minimize carryover. Bound proteins were eluted by adding 50 μL of 2× Laemmli sample buffer followed by boiling. Samples (15 μL) were resolved by SDS–PAGE alongside 2 μL of input sample as a reference.

**Supplementary Data Files**

**Data S1.** GEE labeling data, processed files

**Data S2.** AlphaFold Multimer prediction score and error values for interactions between PU.1 tAD peptides and SMARCD2 SWIFT domain

**Data S3.** Nanoscale HDX-MS raw data

**Data S4.** Mass-spectrometry analyses of HA-SWIFT IP and SMARCA4 (BRG1) IP

**Data S5.** Mass-spectrometry analyses of SWIFT pulldown experiments from cell lysates

**Data S6.** TMT-Mass spectrometry analyses of SMARCD SWIFT pulldown experiments, including SMARCD1/2/3 paralogs, SMARCD2 R290W, and SMARCD2 in the presence of PPSC1 small molecule
